# Neutralization of zoonotic retroviruses by human antibodies: genotype-specific epitopes within the receptor-binding domain from simian foamy virus

**DOI:** 10.1101/2022.11.06.515319

**Authors:** Lasse Toftdal Dynesen, Ignacio Fernandez, Youna Coquin, Manon Delaplace, Thomas Montange, Richard Njouom, Chanceline Bilonga-Ndongo, Felix A. Rey, Antoine Gessain, Marija Backovic, Florence Buseyne

## Abstract

Infection with viruses of animal origin pose a significant threat to human populations. Simian foamy viruses (SFVs) are frequently transmitted to humans, in which they establish a life-long infection, with the persistence of replication-competent virus. However, zoonotic SFVs do not induce severe disease nor are they transmitted between humans. Thus, SFVs represent a model of zoonotic retroviruses that lead to a chronic infection successfully controlled by the human immune system. We previously showed that infected humans develop potent neutralizing antibodies (nAbs). Within the viral envelope (Env), the surface protein (SU) carries a variable region that defines two genotypes, overlaps with the receptor binding domain (RBD), and is the exclusive target of nAbs. However, its antigenic determinants are not understood. Here, we characterized nAbs present in plasma samples from SFV-infected individuals living in Central Africa. Neutralization assays were carried out in the presence of recombinant SU that compete with SU at the surface of viral vector particles. We defined the regions targeted by the nAbs using mutant SU proteins modified at the glycosylation sites, RBD functional subregions, and genotype-specific sequences that present properties of B-cell epitopes. We observed that nAbs target conformational epitopes. We identified three major epitopic regions: the loops at the apex of the RBD, which likely mediate interactions between Env protomers to form Env trimers, a loop located in the vicinity of the heparan binding site, and a region proximal to the highly conserved glycosylation site N8. We provide information on how nAbs specific for each of the two viral genotypes target different epitopes. Two common immune escape mechanisms, sequence variation and glycan shielding, were not observed. We propose a model according to which the neutralization mechanisms rely on the nAbs to block the Env conformational change and/or interfere with binding to susceptible cells. As the SFV RBD is structurally different from known retroviral RBDs, our data provide fundamental knowledge on the structural basis for the inhibition of viruses by nAbs.

## Introduction

Foamy viruses (FVs) are the most ancient of retroviruses [1, 2]. Simian foamy viruses (SFVs) are widespread in nonhuman primates (NHPs), replicate in the buccal cavity, and are transmitted to humans mostly through bites by infected NHPs [2-4]. Such cross-species transmission events currently occur in Asia, Africa, and the Americas. Most individuals known to be infected with zoonotic SFV live in Central Africa, the region at the epicenter of the emergence of HIV-1 and HTLV-1 from their simian reservoirs. In their human hosts, SFVs establish a life-long persistent infection associated with subclinical pathophysiological alterations [5, 6]. Thus far, neither severe disease nor human-to-human transmission have been described, suggesting efficient control of SFV replication and transmission in humans.

SFV infection elicits Env-specific neutralizing antibodies (nAbs) that block the entry of viral particles into susceptible cells *in vitro* [7]. Although they do not block cell-to-cell infection *in vitro*, antibodies prevent cell-associated SFV transmission by transfusion in monkeys [8, 9]. SFV Env is cleaved into three subunits, the leader peptide (LP), the surface subunit (SU), which binds to cells, and the transmembrane (TM) subunit, which carries out fusion (Fig. 1). In several SFV species, there are two variants of the *env* gene, defining two genotypes [10-12]. The two variants differ in a discrete region encoding 250 residues, named SUvar, located at the center of the SU and overlapping with most of the receptor-binding domain (RBD) [13]. The RBD is thus bimorphic and we showed that the SUvar region is the exclusive target of genotype-specific nAbs [7].

**Fig. 1.**
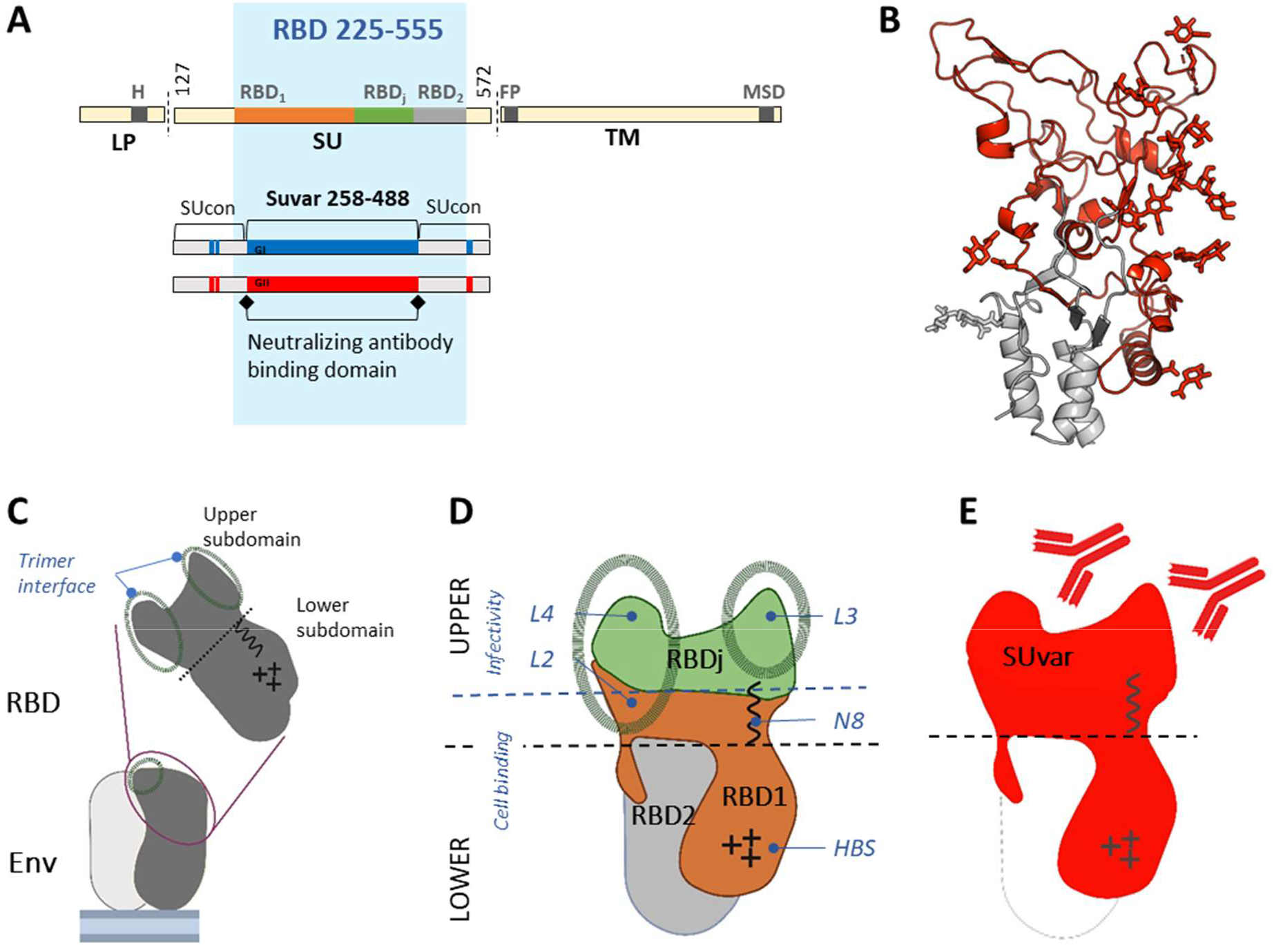
Current knowledge of SFV Env and its RBD. A. Schematic representation of SFV Env. The precursor Env protein is cleaved by furin-like protease(s) at two sites (vertical bars) to generate LP, SU, and TM. The dark sections highlight the transmembrane regions of LP (H) and TM (MSD), and the fusion peptide (F). The minimal continuous RBD (aa 225-555, blue background) comprises two regions essential for SU binding to cells, RBD1 (aa 225-396) and RBD2 (aa 484-555) [13]. The intervening region (aa 397-483), named RBDj, can be deleted without abrogating binding to susceptible cells. Genetic studies on SFV strains circulating in Central Africa have identified two genotypes that differ in the sequence encoding the central part of the SU (SUvar, aa 248-488) domain [11, 12]. SUvar partially overlaps with the RBD and is the exclusive target of nAbs [7]. B. The RBD (aa 218-552) structure from a genotype II gorilla SFV strain is shown as a ribbon (PDB code 8AIC), with SUvar in red and SUcon in grey [14]. Side chains of the glycans are shown and were identified on deglycosylated RBD [14]. Structural elements relevant for the present study are indicated. C. Schematic representation of the two RBD subdomains and their location at the top of Env trimers; the drawing highlights the region involved in trimer assembly [14, 15]. D. RBDj is located at the apex of the RBD upper subdomain; RBD1 forms the second part of the upper subdomain, prolonged in the lower subdomain by an arm wrapping around the stem (RBD2), of which the sequence is conserved. Structural features described in [14] are highlighted on the solvent exposed face of the RBD: the loops mediating trimer interaction (loop 1 is on the internal face and thus not depicted), the conserved glycosylation site (N8), and the HBS. E. The SUvar domain forms the upper RBD subdomain and part of the lower subdomain.

We previously determined the RBD structure at high resolution [14] (Fig. 1A and 1B). The RBDs occupy the space at the top of Env trimer [14, 15] (Fig. 1C). Each RBD has two subdomains. The upper subdomain carries protrusions (loops 1 to 4) that mediate interactions between protomers and the conserved glycosylation site (N8) [14]. The lower subdomain carries the heparan sulfate glycosaminoglycan binding site (HBS) [14]. The RBD has been functionally defined as bipartite [13]. The first essential binding region (RBD1) forms half of the upper subdomain and wraps around the second essential binding region (RBD2) to form the lower subdomain. The central region (called RBDj) is dispensable for cell binding, forms the top of the upper domain, and encompasses loops 3 and 4 (Fig. 1D). The genotype-specific SUvar comprises the entire upper RBD subdomain and one third of its lower subdomain, corresponding grossly to the previously defined RBD1.

The genotype-specific SUvar region is targeted by nAbs [7] (Fig. 1E). Here, we determined the epitope sites on SUvar recognized by plasma nAbs from SFV-infected African hunters. These individuals were infected by SFV of gorilla origin through bites [3], and we previously described their infecting strains, *ex vivo* blood target cells, antibody response, and medical status [5-7, 9, 11, 16, 17]. We expressed the SFV SU as a soluble recombinant protein that competes with SU present on viral vector particles for binding to plasma nAbs in a neutralization assay. We defined the SUvar regions targeted by the nAbs using mutant SU proteins modified at the glycosylation sites, RBD functional subregions, and genotype-specific sequences that present properties of B-cell epitopes. This immunological study and that on the structure of RBD [14] were carried out concomitantly; thus we integrated structural information as it became available and proposed modes of action for SFV-specific nAbs.

## Results

### SFV SU recombinant protein competes with the virus for binding to nAbs

We sought to define nAb epitopes by performing neutralization assays in the presence of recombinant SU protein that competes with the SU within Env present at the surface of foamy viral vector (FVV) particles. While wildtype (WT) recombinant SU binds to the nAbs, allowing cell infection to proceed, SU mutants with altered nAb epitopes do not (Fig. 2A). We tested several recombinant Env and SU expression constructs, of which the production level and stability varied according to the genotype: Gorilla genotype II (GII) Env-derived constructs were expressed at high levels, whereas gorilla genotype I (GI) Env-derived counterparts were poorly expressed and aggregated (Supplementary Fig. 1 and 2 A). To increase the yield and stability of the proteins, we appended the immunoglobulin Fc domain, which acts as a secretion carrier, to the C-terminus of SU. Such chimeric proteins - referred to as immunoadhesins – have been used to produce soluble SFV SU [13]. Nonetheless, the gorilla genotype I (GI) SU immunoadhesin was expressed at insufficient levels. However, we readily produced the chimpanzee genotype I (CI) and GII SU immunoadhesins. We therefore studied epitopes recognized by GII-specific nAbs with the GII immunoadhesin (referred to as ^GII^SU for the WT sequence) and those recognized by GI-specific nAbs with the CI immunoadhesin (^CI^SU), as allowed by the frequent cross-neutralization of gorilla and chimpanzee strains belonging to the same genotype [7] and CI and GI SUvar sequence identity of 70% [11].

**Fig. 2.**
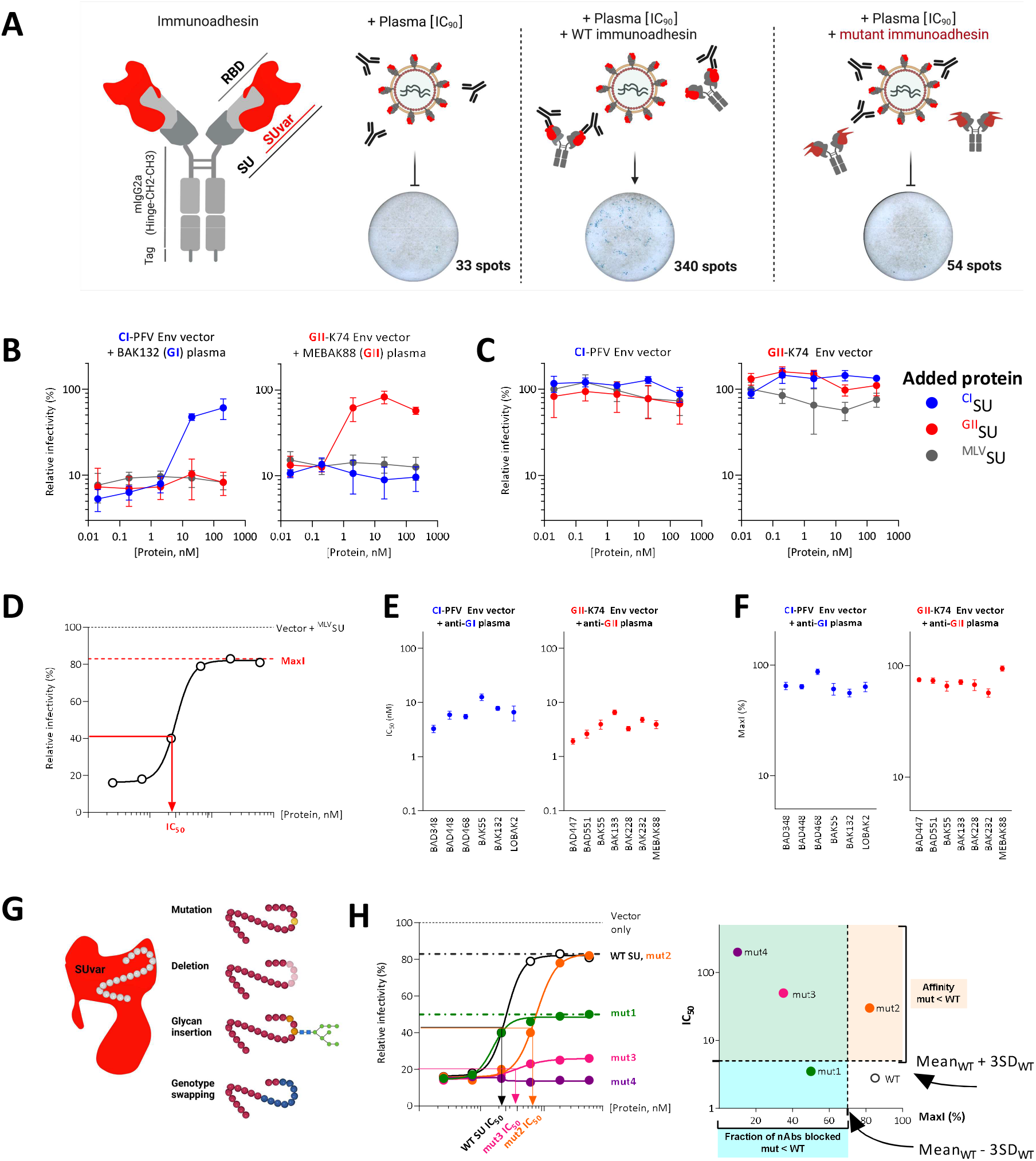
The SFV SU block nAbs. A. Schematic presentation of the neutralization assay using immunoadhesin as competitorSU immunoadhesins are chimeric proteins composed of SU, the constant fragment of murine IgG2a, and a double Strep-tag. The SU, RBD, and SUvar are highlighted in the drawing. Plasma samples were diluted to achieve a reduction in the number of FVV-transduced cells by 90%. WT SU compete with Env on FVV for binding by nAbs, resulting in a higher number of FVV-transduced cells; mutations in SU that affect binding by nAbs result in inefficient competition and reduced FVV transduction. Representative images of wells with FVV-transduced cells stained by X-gal are shown. B. The BAK132 (anti-GI) and MEBAK88 (anti-GII) plasma samples were incubated with SU and the mix added to FVVs expressing matched Env before titration. The relative proportion of transduced cells is expressed as the percentage of cells transduced by untreated FVVs (no plasma and no protein), is referred to as the relative infectivity, and is presented as a function of protein concentration. The addition of ^CI^SU (blue symbols) inhibited the action of nAbs from sample BAK132, as shown by increased CI-PFV Env FVV relative infectivity, whereas ^GII^SU (red symbols) had no effect. Conversely, ^GII^SU inhibited the action of nAbs from sample MEBAK88. ^MLV^SU (grey symbols) had no effect on the plasma antibodies. C. The infectivity of the CI-PFV and GII-K74 Env vectors was quantified in the presence of ^CI^SU, ^GII^SU, and ^MLV^SU; the relative infectivity is presented as a function of protein concentration. D. Schematic representation of a WT SU titration curve, summarized by two parameters, MaxI and IC_50_. E and F. Thirteen pairs of plasma samples and genotype-matched SU were tested at least five times for their activity against FFVs. The mean and standard error of the mean of the IC_50_ (panel E) and MaxI (panel F) are shown for the CI-PFV Env vectors and anti-GI plasma samples (blue symbols) and the GII-K74 Env vectors and anti-GII plasma samples (red symbols). G. To define neutralizing epitopes, we synthesized SU mutants with four types of alterations in SUvar: mutations, deletion, glycan insertion, and swapping of sequences with the second genotype. H. Schematic representation of titration curves corresponding to SU mutants that lose their capacity to block a fraction of nAbs (mut1, green), block all nAbs with reduced affinity (mut2, orange), or block a fraction of nAbs with reduced affinity (mut 3, pink) or those with no blocking activity (mut4, purple). Two parameters were defined using the curves, MaxI and IC_50_, and were compared to those obtained with WT SU (panels E and F) to detect significant differences in binding (see Materials and Methods).

We validated the competition strategy with genotype-matched and mismatched immunoadhesins (Fig. 2B, Supplementary Tables 1 and 2, Supplementary Fig. 3). We tested plasma samples from gorilla SFV-infected hunters at the dilution required for 90% reduction of FVV infectivity (IC_90_) to allow Env-specific antibody saturation by the competitor SU. Thus, plasma samples were tested at the same nAb concentration and responses from different individuals could be compared. We incubated SU with plasma samples before addition to a FVV expressing the matching Env protein and titration of residual infectivity. Both ^CI^SU and ^GII^SU blocked the action of nAbs present in genotype-matched plasma samples in a concentration-dependent manner but failed to block the activity of genotype-mismatched samples (Fig. 2B). Importantly, the immunoadhesins did not affect SFV entry (Fig. 2C, [18]), allowing their use as competitors for binding to nAbs in an infectivity assay. The unrelated MLV SU construct (^MLV^SU) had no impact on either the neutralizing activity of plasma samples or SFV infectivity (Fig. 2B and 2C).

We noticed that immunodhesins did not fully block plasma nAbs and hypothesized that this could be due to their monomeric state and lack of the epitopes formed by quaternary structures. In addition, heterogenous glycosylation can lead to partial resistance to neutralization [19-22]. Thus, we evaluated the impact of SU oligomerization and mammalian-type glycosylation on the capacity to block nAbs by testing the following constructs: SU monomers produced in mammalian and insect cells, trimeric Env ectodomains produced in insect cells, and dimers of monomeric SU (i.e., immunoadhesin) produced in mammalian cells. Since all constructs had the same capacity to block the nAbs (Supplementary Fig. 2 B), we used the only recombinant proteins available for both genotypes, the SU immunoadhesins.

We titrated 11 samples against matched ^CI^SU or ^GII^SU, and one sample from a coinfected individual against both SUs (Supplementary Table 2). The effect of the SU on nAbs can be described by two sample-dependent parameters (Fig. 2D). The first parameter is the affinity, quantified by the protein concentration required to inhibit 50% of the neutralizing activity (IC_50_). IC_50_ values ranged between 3.3 and 12.6 nM for ^CI^SU and 2.9 and 12.5 nM for ^GII^SU, depending on the plasma sample (Fig. 2E). The second parameter is the fraction of neutralizing activity that can be blocked by the SU. Plasma samples contain a polyclonal population of nAbs, some of which bind monomeric SU in its immunoadhesin format, whereas others may not. Taking FVV treated with ^MLV^SU without plasma as a reference value for full infectivity (100%), ^CI^SU and ^GII^SU restored the infectivity of plasma-treated FVV to between 56% and 94% of infectivity. This value is referred to as maximum infectivity (MaxI) and represents the fraction of nAbs blocked by the SU (Fig. 2D and 2F). We then designed panels of mutant SU proteins (Fig. 2 G) and compared their capacity to block nAbs to that of WT SU based on changes in their IC_50_ and MaxI. We first studied GII-specific nAbs using ^GII^SU before testing GI-specific nAbs using ^CI^SU.

### A fraction of nAbs targets glyco-epitopes

The GII SUvar domain carries seven of the 11 SU glycosylation sites (Fig. 3A and 3B). We assessed whether these glycans are recognized by nAbs or whether they may shield epitopes from nAb recognition. We produced ^GII^SU in mammalian cells in the presence of the mannosidase inhibitor kifunensine, which prevents complex-type glycan addition, and treated a fraction with endo-H to remove all glycans. The absence of complex glycans had no impact on nAb blockade by ^GII^SU (Fig. 3C), in accordance with the data obtained with GII Env constructs expressed in insect cells, which also lack complex glycans (Supplementary Fig. 2). By contrast, the removal of all glycans by Endo-H decreased nAb blockade and protein titration showed lower affinity for the nAbs present in three of four plasma samples (Fig. 3D). We verified that this was not due to aggregation induced by Endo-H treatment (Supplementary Fig. 4). The fact that the removal of glycans did not enhance nAb blockade in the samples tested indicates that the glycans do not shield the majority of nAb epitopes on SFV Env.

**Fig. 3.**
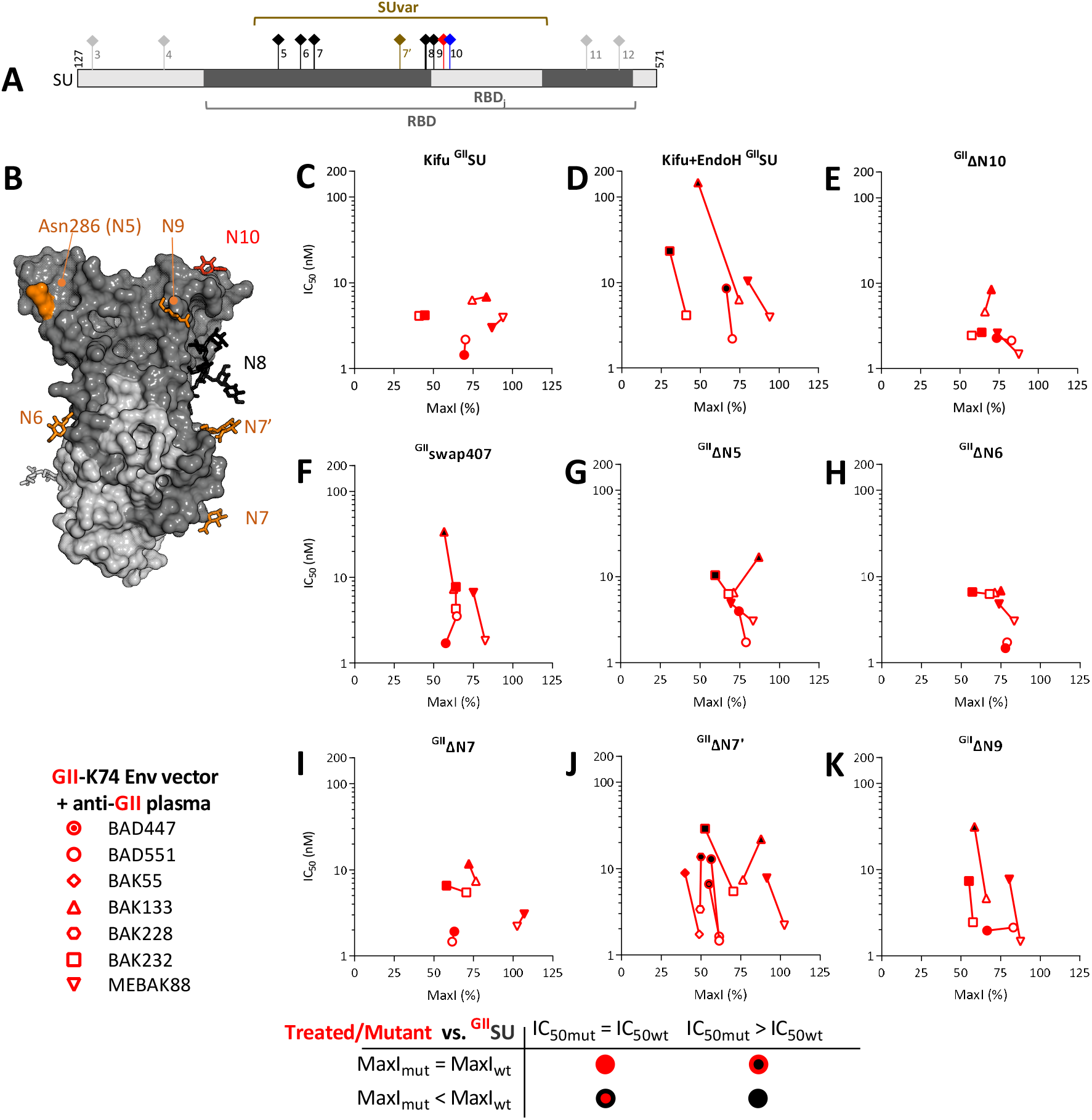
SFV-specific nAbs recognize glycans on SUvar. A. Schematic representation of N-glycosylation sites on the SU.The N7’ site (position 374, brown symbol) is absent from CI-PFV but present in zoonotic gorilla SFV and several chimpanzee SFVs [11]. The N8 site (position 391, bold stem) is strictly conserved and required for SU expression [23]. The N10 site has a genotype-specific location (N411 in GII strains, red symbol; N422 or 423 in GI/CI strains, blue symbol). The glycosylation sites outside SUvar are shown in grey. B. The RBD is shown as a solvent accessible surface, with SUvar in dark grey and SUcon in light grey. The glycans resolved in the SFV RBD X-ray structure are shown as sticks, colored in orange for those for which a deletion mutant was constructed, in black for the N8 and in grey for that located on SUcon; the N5 glycan was poorly resolved and N286 is colored to indicate the localization of its anchor. C and D. To determine whether SFV-specific nAbs target residue glycosylated epitopes, vectors carrying GII-K74 Env were mixed with four genotype-matched plasma samples previously incubated with untreated, kifunensine treated (C), or kifunensine and endo-H treated (D) ^GII^SU at several concentrations. E. To test whether nAbs target the genotype-specific N10 glycosylation site, the SU in which N10 was removed (^GII^ΔN10) was incubated with four genotype-matched plasma samples. F. Residues 407-413 were swapped with those from the GI-D468 strain and the resulting ^GII^swap407 was tested for its ability to block four anti-GII plasma samples. The glycosylation sites located on the SUvar were inactivated one by one (except N8) and tested for their inability to block four GII-specific plasma samples (G to K). ^GII^ΔN7’ was tested against three additional samples to confirm its impact on nAbs (J). For each plasma sample, the IC_50_ is presented as a function of MaxI for untreated and enzyme-treated ^GII^SU (C and D) or ^GII^SU and mutants (E to K). The IC_50_ and MaxI values for untreated ^GII^SU are presented as open symbols and are from the same experiment in which the mutant SU were tested. For the enzyme-treated and mutant SU, the symbols are colored according to the IC_50_ and MaxI thresholds that were used to statistically define significant differences from ^GII^SU.

The N10 glycosylation site is located at different positions for the two genotypes (N422/N423 on GI-D468/CI-PFV strains, N411 on the GII-K74 strain) and we hypothesized that it may be part of a potential epitopic region. The N10 glycosylation knock-down mutant (^GII^ΔN10) was as efficient as ^GII^SU in blocking nAbs from all plasma samples (Fig. 3E). N10 belongs to a stretch of seven genotype-specific residues that were replaced by those from the GI-D468 strain. The chimeric protein (^GII^swap407) blocked nAbs as efficiently as ^GII^SU for three samples and showed decreased affinity against nAbs for one sample (Fig. 3F). Thus, GII-specific nAbs occasionally target epitopes affected by the N10 glycosylation site.

To identify other glycosylation sites targeted by nAbs, we knocked them down one by one, except for N8, which is critical for SU expression [23]. The ^GII^ΔN7’ mutant showed reduced capacity to block nAbs from five of the seven samples (Fig. 3J). ^GII^ΔN5 and ^GII^ΔN9 showed reduced capacity to block nAbs from one or two plasma samples (Fig. 3G and 3K). Knock down of N6 and N7 had no impact (Fig. 3H and 3I). Overall, our data indicate that most glycans are not part of epitopes, except those at the N7’ glycosylation site.

### SFV-specific nAbs target the upper RBD subdomain

Next, we aimed to locate nAb epitopes on the SU subdomains. On the upper domain, RBDj could be deleted without fully compromising cell binding, whereas the loops it harbors (Fig. 1, 4A and 4B) are essential for infectivity [13, 14]. We therefore constructed a mutant SU with RBDj deleted. ^GII^ΔRBDj was highly impaired in blocking nAbs from three individuals: the IC_50_ values were above the highest concentration tested (200 nM) and the MaxI values were close to 10%, which is the value of plasma samples incubated with the irrelevant ^MLV^SU (Fig. 4C). By contrast, ^GII^ΔRBDj blocked nAbs from the BAD551 plasma sample as efficiently as ^GII^SU.

**Fig. 4.**
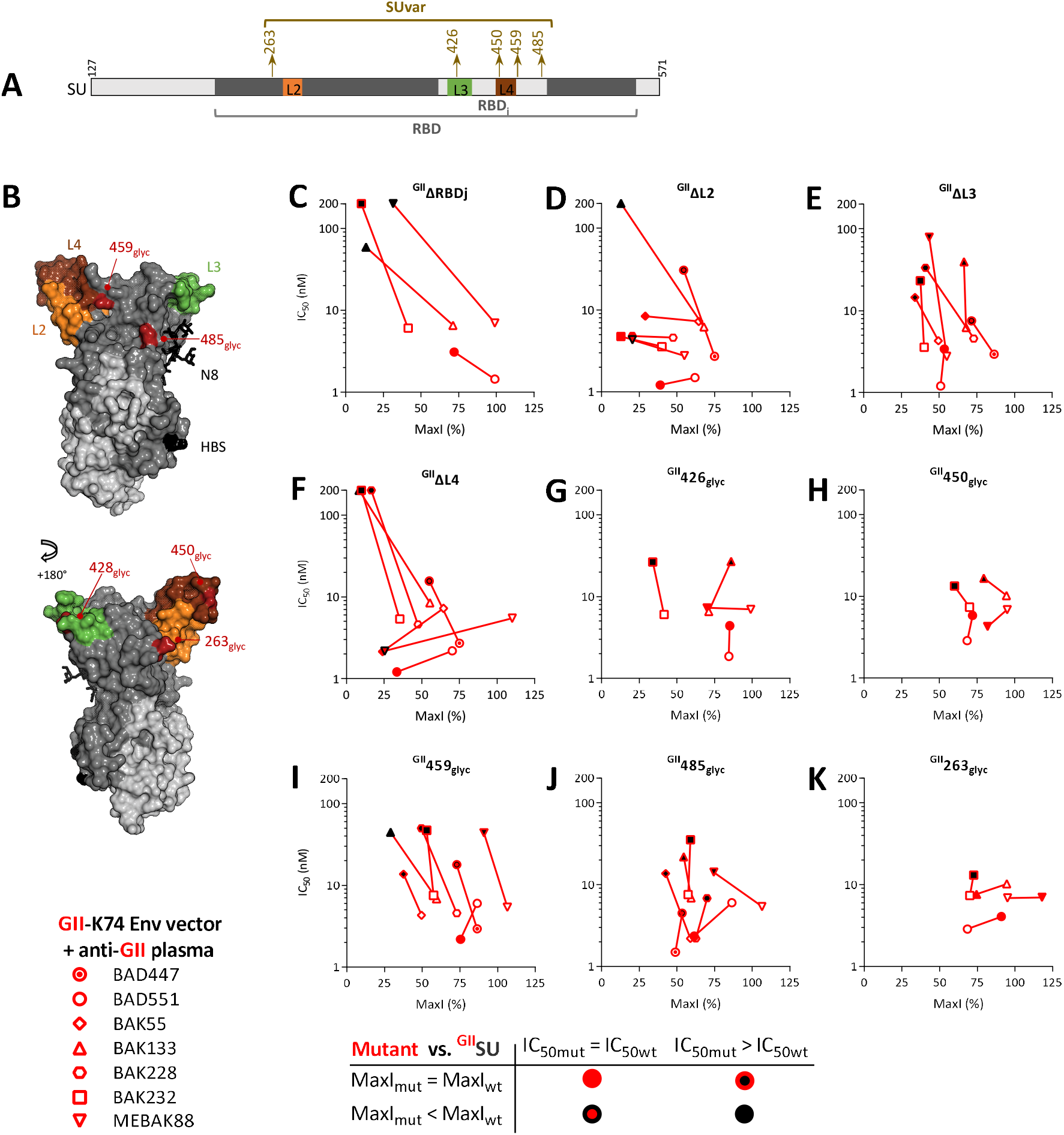
Most plasma samples contain nAbs that recognize the RBDj domain. A.Schematic representation of the SU, with the RBD, the loops, and the inserted glycosylation sites. B. The RBD is shown as a solvent accessible surface with SUvar in dark grey, SUcon in light grey, the side chain from the N8 glycans in dark grey, L2 in orange, L3 in green, L4 in maroon, HBS in black, and the positions of glycosylation site insertions in dark red. To locate SFV-specific nAb epitopes on the upper RBD subdomain, ^GII^SU and mutants with RBDj deleted (^GII^ΔRBDj, panel C), deleted loops (^GII^ΔL2, ^GII^ΔL3, and ^GII^ΔL4 in panels D-F), and glycans inserted in putative epitopes (at positions 426, 450, 459, 485, and 263 in panels G to K) were tested against at least four plasma samples. Those for which the capacity to block nAbs was the most altered were then tested on additional samples. For each plasma sample, the IC_50_ is presented as a function of MaxI for ^GII^SU and the mutant SU. The IC_50_ and MaxI values for ^GII^SU are presented as open symbols and are from the same experiment in which the mutant SU were tested. Symbols are colored according to the IC_50_ and MaxI thresholds used to define statistically significant differences from ^GII^SU.

Among the four loops that emanate from the upper domain L2 (276-281), L3 (416-436), and L4 (446-453) are exposed to solvent and mobile and therefore considered to be candidate epitopic regions [14]. We deleted the loops individually (Fig. 4D-F). ^GII^ΔL3 blocked all GII-specific samples but one, although with lower affinity. ^GII^ΔL2 and ^GII^ΔL4 showed plasma-dependent effects, with three patterns: (1) full activity (such as against BAD551) (2), no or strongly reduced of activity (for example, against BAK133), (3) or a reduced MaxI with unchanged affinity (such as MEBAK88). The last pattern probably reflects the presence of nAbs targeting different epitopes within the same plasma sample, some epitopes being altered on mutant SU, whereas others are not.

Before the 3D structure was available, we relied on the insertion of N-linked glycosylation sequences (NXS/T) to disrupt potential epitopes. We selected six genotype-specific sequences predicted to have a disordered secondary structure or to contain a B-cell epitope and built mutant SU with a glycosylation site inserted in the targeted sequence (Supplementary Table 3). Five of the glycosylation sites were inserted in the upper subdomain, with three located within or close to L3 and L4 loops, resulting in decreased affinity against certain plasma nAbs: ^GII^426_glyc_ in L3, ^GII^450_glyc_ in L4, ^GII^459_glyc_ after L4 (Fig. 4G-I). The fourth mutant with reduced affinity against most plasma nAbs, ^GII^485_glyc_, is located in RBDj in the vicinity of N8, which plays a major structural role [14] (Fig. 4J). Finally, ^GII^263_glyc_ had no or only a modest effect on nAb activity, which is consistent with its location at the center of the trimer, which is likely not accessible to antibodies (Fig. 4K, [14]). Overall, glycan insertions confirmed the frequent recognition of RBDj by nAbs.

### SFV-specific nAbs target epitopes in the lower RBD subdomain

In the lower subdomain, the HBS was a candidate epitopic region (Fig. 5). The GII Env ectodomain mutated on HBS residues (^GII^K342A/R343A or ^GII^R356A/R369A, [14]) blocked nAbs from three samples as efficiently as their WT counterpart and showed lower affinity for the MEBAK88 plasma sample (Fig. 5C and 5D). Thus, nAb activity is modestly affected by mutated HBS residues.

**Fig. 5.**
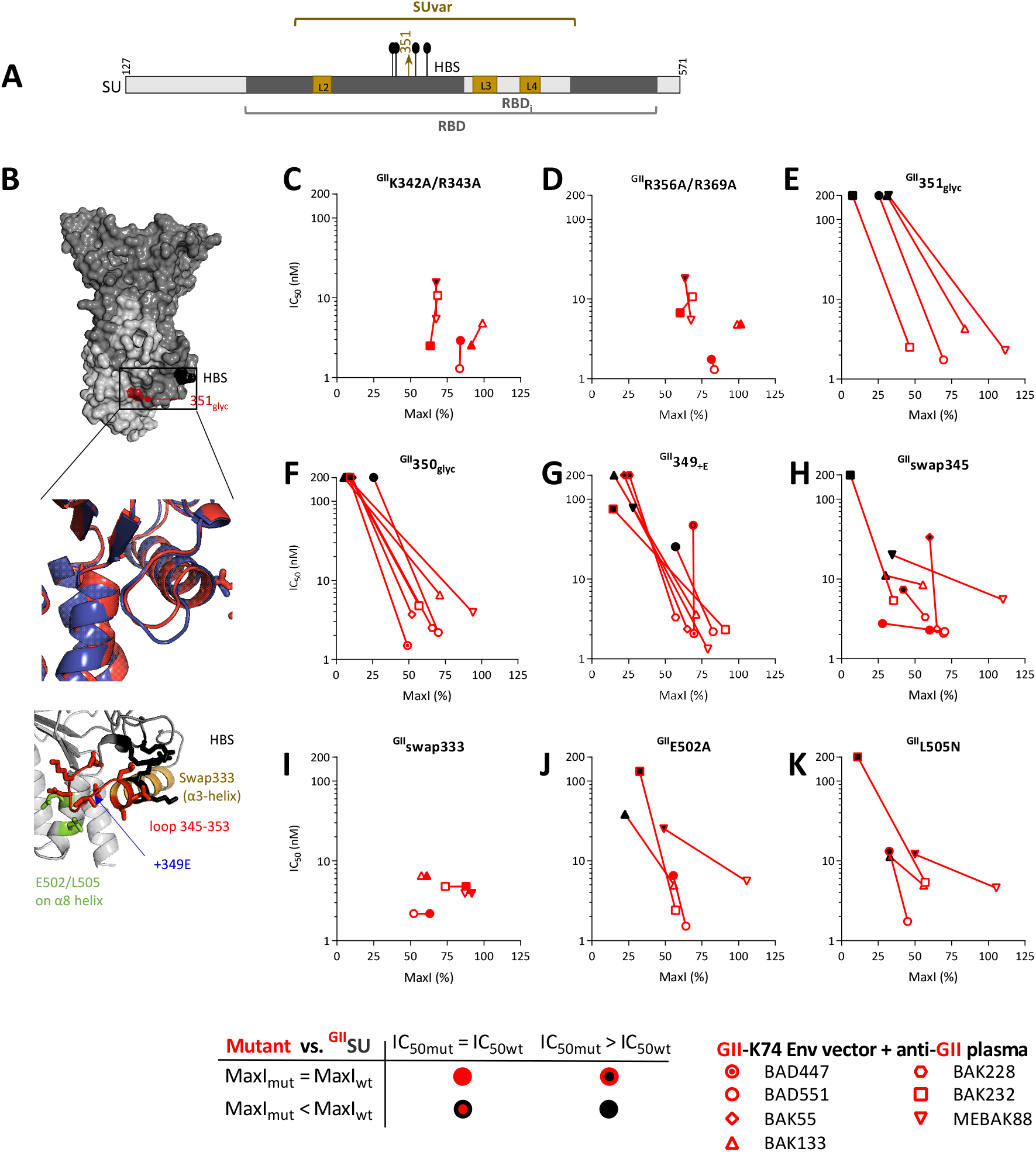
Epitope disruption by glycan insertion reveals an epitope site on the lower RBD subdomain. A. Schematic representation of the SU, with the RBD, loops, HBS, and site of glycan insertion at position 351 on SU. B. The RBD is shown as a solvent accessible surface with SUvar in dark grey, SUcon in light grey, HBS in black, and the glycan insertion site in dark red. In the top insert, the 345-353 loop and adjacent helix are presented as ribbons and side chains for GII-K74 (red) and CI-PFV (blue, AlphaFold2-predicted structure). In the bottom insert, the 345-353 loop (red) and adjacent helix are presented as ribbons and side chains for GII-K74, with colored residues indicating the HBS (black), swap333 construct (light brown), insertion site at position 349, and two residues on the adjacent helix (green). To locate SFV-specific nAb epitopes on SU functional domains, vectors carrying GII-K74 Env were treated with GII-specific plasma samples previously incubated with the Env ectodomain carrying WT or mutated HBS (^GII^K342A/R343A in panel C and ^GII^K356A/R369A in panel D). E. A candidate genotype-specific sequence(Supplementary Table 1) was disrupted by inserting glycan in the SU at position 351 and tested for its capacity to block nAbs. F to K. Five mutant SU were tested to characterize the epitopic region in the 345-353 loop. All mutants were tested against at least four plasma samples. Those for which the capacity to block nAbs was the most altered were then tested on additional samples. For each plasma sample, the IC_50_ is presented as a function of MaxI for ^GII^SU and the mutant SUs. The IC_50_ and MaxI values of ^GII^SU are presented as open symbols and are those from the same experiment in which the mutants were tested. Symbols are colored according to the IC_50_ and MaxI thresholds used to statistically define significant differences from ^GII^SU. We applied the same threshold values for the analyses of the Env ectodomain, which had the same inhibitory activity as ^GII^SU (Supplementary Fig. 2). ^GII^swap333 was tested twice at three concentrations and showed similar blocking capacity as ^GII^SU; the IC_50_ values were arbitrarily set to the same level as those of ^GII^SU (see Materials and Methods).

In the SU series with glycan insertions in predicted epitopes (Supplementary Table 3), one had a glycan inserted in the lower RBD subdomain (^GII^351_glyc_) and showed a strongly reduced capacity to block the nAbs (Fig. 5E). The modified residue is located on a solvent-exposed loop (residues 345-353) at the base of the SUvar and is close to the HBS (Fig. 5B). ^GII^351_glyc_ was the first mutant to indicate that nAbs recognize a region of the RBD to which no function has yet been attributed. We therefore focused the following experiments on this novel putative epitope.

As ^GII^351_glyc_ was prone to aggregation (Supplementary Fig. 4H), we produced a second batch, half of which was purified by affinity chromatography (standard protocol) and the other half further purified by size exclusion chromatography (SEC) to eliminate the aggregates (Supplementary Fig. 5A). Both ^GII^351_glyc_ preparations were unable to block nAbs from the plasma samples (Supplementary Fig. 5B-E). Glycan insertion at an adjacent position in ^GII^350_glyc_ led to the loss of nAb blockade and reduced aggregation (Fig. 5F, Supplementary Fig. 4H). The GII 345-353 loop is one residue shorter than that of the GI-D468 and CI-PFV strains (Fig. 5B). The mutant SU with an extra glutamic acid at position 349 (^GII^ 349_+E_) showed a strongly reduced capacity to block the nAbs (Fig. 5G). We replaced the seven GII residues by the eight residues from the GI-D468 strain; ^GII^swap345 showed a reduced ability to block nAbs from five of the seven plasma samples (Fig. 5H). By contrast, swapping the α3-helix located N-terminal to the loop (^GII^swap333) had no impact on the capacity to block the nAbs (Fig. 5I). Finally, we tested the substitution of residues on the α8-helix facing the 345-353 loop; ^GII^E502A and ^GII^L505N showed a reduced capacity to block the nAbs (Fig. 5K and 5K). Overall, these experiments define the 345-353 loop and adjacent α8-helix as a major epitopic region on GII RBD.

We directly tested the capacity of plasma nAbs to block the entry of FVVs expressing Env mutated in the 345-353 loop to confirm the data obtained with the competition strategy. We chose to express a mutated Env corresponding to the ^GII^swap345 mutant, as its cell binding capacity is unaltered in the immunoadhesin format (see below). FVVs with a swapped 345-353 Env loop were produced at the same level as those expressing WT GII-K74 Env and had similar infectious titers and cell binding capacity (Fig. 6A to 6C). We carried out neutralization assays for both vectors using the seven GII-specific plasma samples. In its soluble form, the ^GII^swap345 protein blocked the neutralizing activity from five plasma samples less efficiently than the ^GII^SU protein (Fig. 5H). FVVs carrying ^GII^swap345 SU were less susceptible than those carrying ^GII^SU to plasma nAbs from three individuals (BAK133, BAK232, BAK228), as reflected by at least two-fold lower neutralization titers against the ^GII^swap345 carrying vector (Fig. 6G to 6I). Plasma samples from individuals BAK55 and MEBAK88 showed similar neutralization titers against both vectors (Fig. 6F and 6J). The ^GII^swap345 protein blocked nAbs from two plasma samples as efficiently as the ^GII^SU protein: the sample from individual BAD551 showed similar neutralizing titers against both vectors, whereas the sample from individual BAD447 neutralized the ^GII^swap345-carrying vector more efficiently than that carrying ^GII^SU (Fig. 6D and 6E). Thus, despite an imperfect match between the competition assay and direct neutralization data, testing epitope modification in the context of viral particles confirmed that loop 345-353 is recognized by plasma nAbs.

**Fig. 6.**
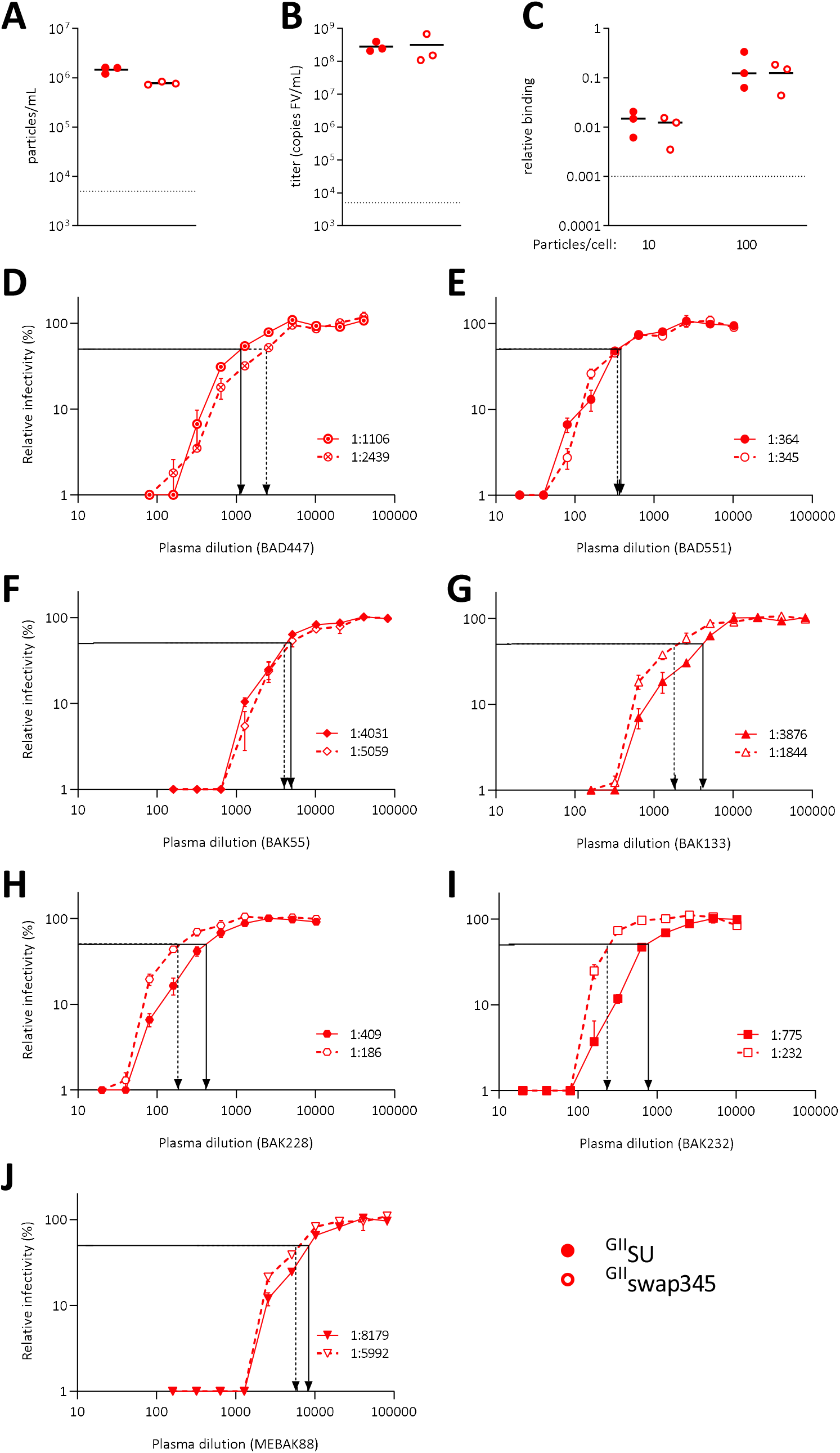
The 345-353 loop is targeted by nAbs at the surface of viral vector particles. A. Three batches of FVVs carrying Env with ^GII^SU or ^GII^SUswap345 were produced. The concentration of vector particles was quantified by RT-qPCR amplification of the *bgal* transgene. B. FVV infectious titers were quantified on BHK-21 cells. C. FVVs carrying WT and mutated Env were incubated with HT1080 cells at 10 or 100 particles/cells on ice for 1 h before washing and quantification of *bgal* mRNA incorporation in the vector particles and the *gapdh* gene of susceptible cells. The levels of *bgal* and cellular *gapdh* mRNA were quantified by RT-qPCR; the ΔΔCt method was used to calculate the relative number of FVV particles bound to cells. The dotted lines in panels A to C represent the quantification threshold and the black lines the mean values from the three FVV batches. The infectious titers, particle concentration, and levels of bound particles from FVVs carrying mutant or WT SU were compared using the paired t test and all p values were > 0.05. D to J. Neutralization assays were carried out by transducing BHK-21 cells with SFV vectors carrying WT (plain symbols) or swap345 (open symbols) GII Env. Vectors were previously incubated with 10 serial dilutions of human plasma samples; the lowest dilution ranged between 1:20 or 1:160 according to the donors’ neutralization titers [7]. Assays were performed twice in triplicate and the results from one experiment are shown. Cells were transduced with untreated FFV to provide the reference value. Relative infectivity was calculated for wells treated with plasma samples and is expressed as the percentage of the reference value. Relative infectivity (mean, SEM) is presented as a function of the plasma dilution. Arrows indicate neutralization titers against vectors carrying WT (plain line) and swap345 (broken line) GII Env. Plasma samples are the same as those presented in Fig. 3 to 5: D, BAD447; E, BAD551; F, BAK55; G, BAK133; H, BAK228; I, BAK232; and J, MEBAK88.

### GI and GII-specific nAbs target different epitopes

We applied a similar strategy using GI-specific plasma samples and ^CI^SU harboring the mutations with the greatest impact on GII-specific epitopes. We present these data, including a comparison with those obtained with the GII-specific samples (Fig. 7, blue symbols (GI) over light red lines (GII)). GI-specific nAbs targeted glycans removed by Endo-H but not complex glycans and did not recognize N10, for which the location is genotype-specific (Fig. 7A-7C). We could not test the recognition of N7’, as it is absent from the CI-PFV strain.

**Fig. 7.**
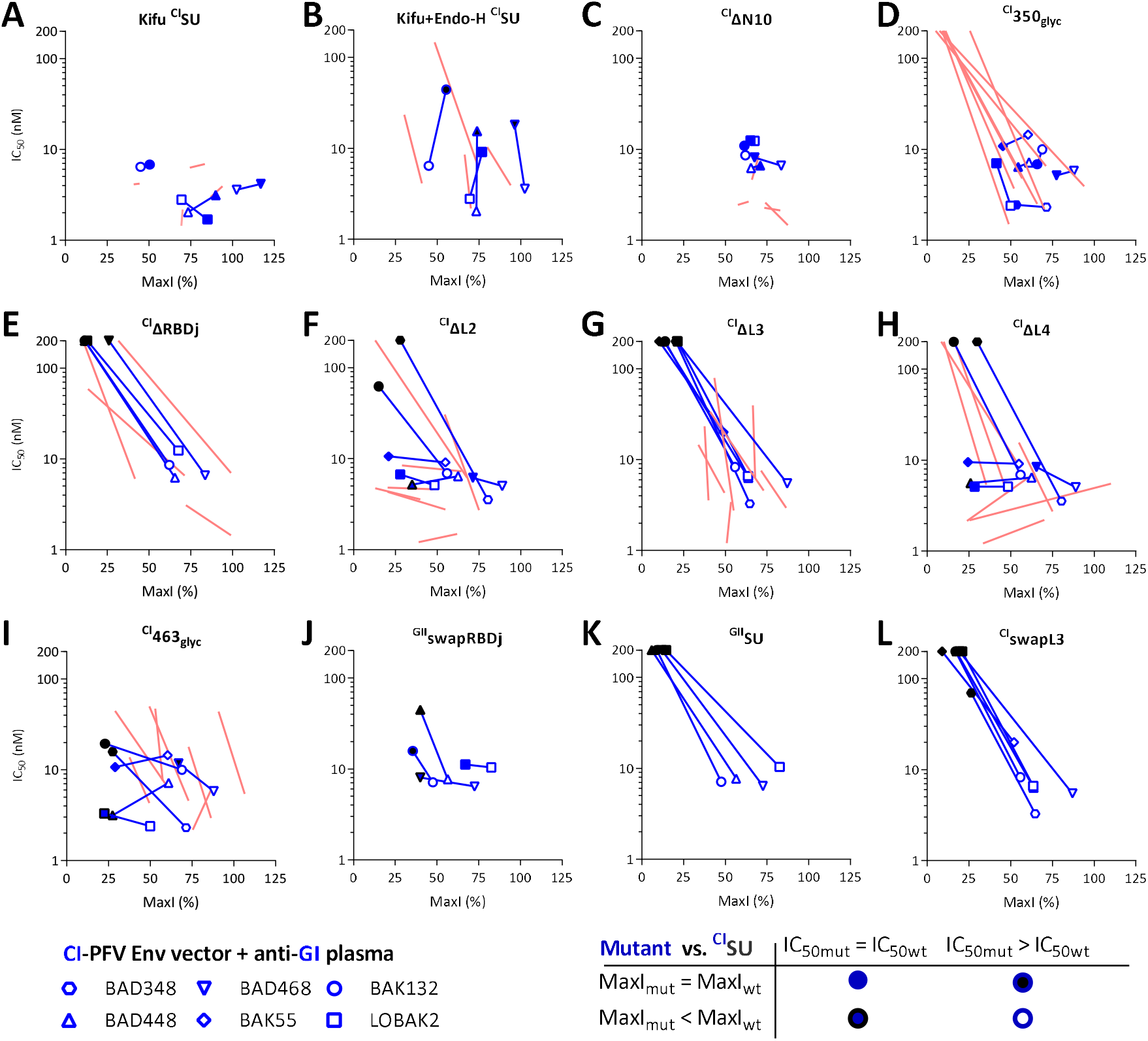
GI and GII-specific nAbs target different epitopesCI SU with mutations matching the most informative GII SU mutations were tested for their capacity to block nAbs from GI-specific plasma samples. A, Kifunensin-treated ^CI^SU; B, Kifunensin and endoH-treated ^CI^SU; C, ^CI^ΔN10; D, ^CI^350_glyc_; E, ^CI^ΔRBDj; F, ^CI^ΔL2; G, ^CI^ΔL3; H, ^CI^ΔL4; I, ^CI^463_glyc_; J, ^GII^swapRBDj; K, ^GII^SU; L, ^CI^swapL3. All mutants were tested against four plasma samples. Those for which the capacity to block nAbs was the most altered were then tested on additional samples. For each plasma sample, the IC_50_ is presented as a function of MaxI for the ^CI^SU and mutant SU. The IC_50_ and MaxI values of ^CI^SU are presented as open symbols and are those from the same experiment in which the mutant SU were tested. For mutant SU, the symbols are colored according to the IC_50_ and MaxI thresholds used to statistically define significant differences from ^CI^SU. The red lines correspond to data obtained with GII-specific plasma samples against equivalent constructs (Panels 3C, 3D, 3E, 5F, 4C, 4D, 4E, 4F, and 4I match panels 7A to 7H, respectively).

The most striking difference between GI and GII epitopes was around residue 350: in sharp contrast to ^GII^350_glyc_ (Fig. 6F), ^CI^350_glyc_ fully blocked GI-specific nAbs (Fig. 7D). In addition, the two chimeric GII SUs with GI residues at positions 333 to 345 and 345 to 351 did not block anti-GI nAbs (Supplementary Fig. 6). Neither ^CI^ΔRBDj nor ^CI^ΔL3 proteins blocked the nAbs (Fig. 7E and 7G). Thus, L3 is more important for GI-than GII-specific nAbs. Indeed, ^GII^ΔL3 blocked GII-specific nAbs, although with lower affinity than ^GII^SU (Fig. 4 E). ^CI^ΔL2 and ^CI^ΔL4 showed sample-dependent effects, as their GII counterparts (Fig. 7F and 7H). ^CI^463_glyc_ blocked GI-specific nAbs, with a reduced MaxI (Fig. 7I), indicating that a fraction of GI-specific nAbs was not blocked, whereas the other was blocked as efficiently as by ^CI^SU. The corresponding ^GII^459_glyc_ mutant blocked GII-specific nAbs, but with lower affinity than ^GII^SU (Fig. 4I). Overall, nAbs from GI-and GII-specific plasma samples target different epitopes.

The mutations on the SU may have induced local conformational changes with epitope-specific effects or global changes with nonspecific effects. The RBDj deletion had a likely epitope-specific effect on the GII backbone, as ^GII^ΔRBDj retained its capacity to block one sample containing nAbs that preferentially targeted epitopes outside the RBDj, indicating that a part of the protein was correctly folded (Fig. 4C). However, ^CI^ΔRBDj and ^CI^ΔL3 did not display any nAb blocking activity. Thus, we confirmed the results using chimeric SU: ^GII^swapRBDj (i.e., GI RBDj in ^GII^SU) blocked nAbs from the four anti-GI plasma samples with a similar or modestly reduced capacity relative to that of ^CI^SU (Fig. 7J), whereas ^GII^SU had no blocking activity (Fig. 7K). ^CI^swapL3 (GII L3 in ^CI^SU) did not block anti-GI nAbs (Fig. 7L). These data suggest that either GI-specific nAbs recognize the L3 loop.

### Human plasma samples do not target linear epitopes on SUvar

We sought to map additional epitopes by screening plasma samples for binding to synthetic peptides. None of eight plasma samples recognized synthetic peptides spanning loops on the upper domain (Fig. 8), indicating that they may be part of conformational epitopes. Furthermore, we screened 17 plasma samples for binding to 37 synthetic peptides spanning the SUvar domain from three viral strains (Supplementary Tables 4 and 5). By ELISA, seven of 17 (41%) plasma samples from SFV-infected individuals reacted against at least one peptide; five locations represented by peptides with GI and/or GII sequences were recognized by at least one plasma sample (Supplementary Fig. 7). Plasma antibodies bound to GI and/or GII peptides irrespective of the SFV genotype against which they were raised. Overall, the peptide binding study showed that most SFV-specific antibodies do not recognize genotype-specific linear epitopes.

**Fig. 8.**
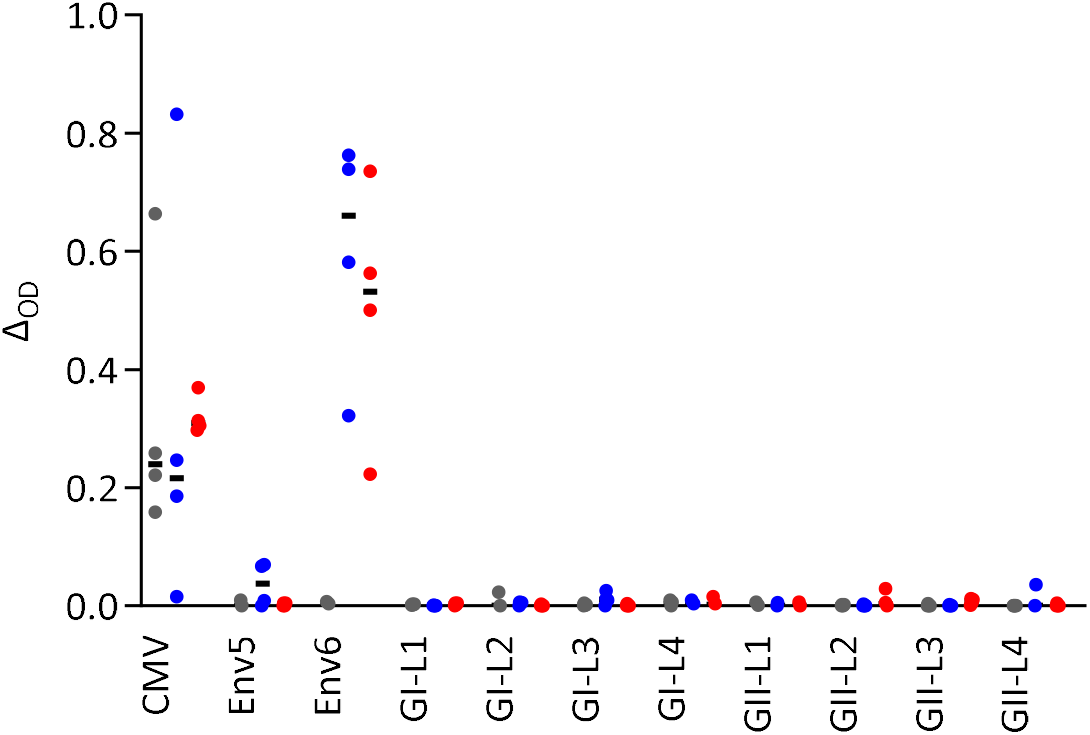
Plasma antibodies do not bind to peptides covering the loops from the upper RBD subdomain. Twelve plasma samples from African hunters (Supplementary Table 4) were tested for binding to peptides overlapping loops located in the upper subdomain of the RBD (Supplementary Table 5).Plasma samples from four uninfected (grey symbols), four GI-infected (blue symbols), and four GII-infected (red symbols) individuals were tested. The CMV and SFV Env6 peptides were used as positive controls and the SFV Env5 peptide as a negative control [24]. The peptide diluent was used as the negative control and antibody binding to peptides is expressed as the difference in OD (Δ_OD_ = OD_test_ – OD_control_). The responses are presented as Δ_OD_ (y-axis) for each peptide (x-axis).

### Genotype-specific residues are involved in SU protein binding to susceptible cells

All SU used for epitope mapping were tested for their capacity to bind to human HT1080 cells, which are highly susceptible to gorilla and chimpanzee SFVs [16] (Fig. 9, Supplementary Fig. 8). SU binding was enhanced after glycan removal, possibly through reduced steric hindrance. Deletion of the RBDj reduced binding to susceptible cells, while one-by-one deletion of loops L2, L3, and L4 abolished it. We observed genotype-specific differences; the RBDj deletion had a stronger impact on GII than on CI SU (6-vs. 2.5-fold reduction). Conversely, glycan insertion after L4 had only a moderate impact on the GII SU (^GII^459_glyc_, 6-fold reduction) but abolished binding of the CI SU (^CI^463^glyc^, ≈100-fold reduction). In the 345-353 loop, glycan insertion strongly affected the binding of GII SU (^GII^350_glyc_ and ^GII^351_glyc_, ≈50-fold reduction), but there was no effect on ^CI^SU, mirroring the recognition by the nAbs. Overall, our data support that certain residues involved in cell binding may be genotype-specific.

**Fig. 9.**
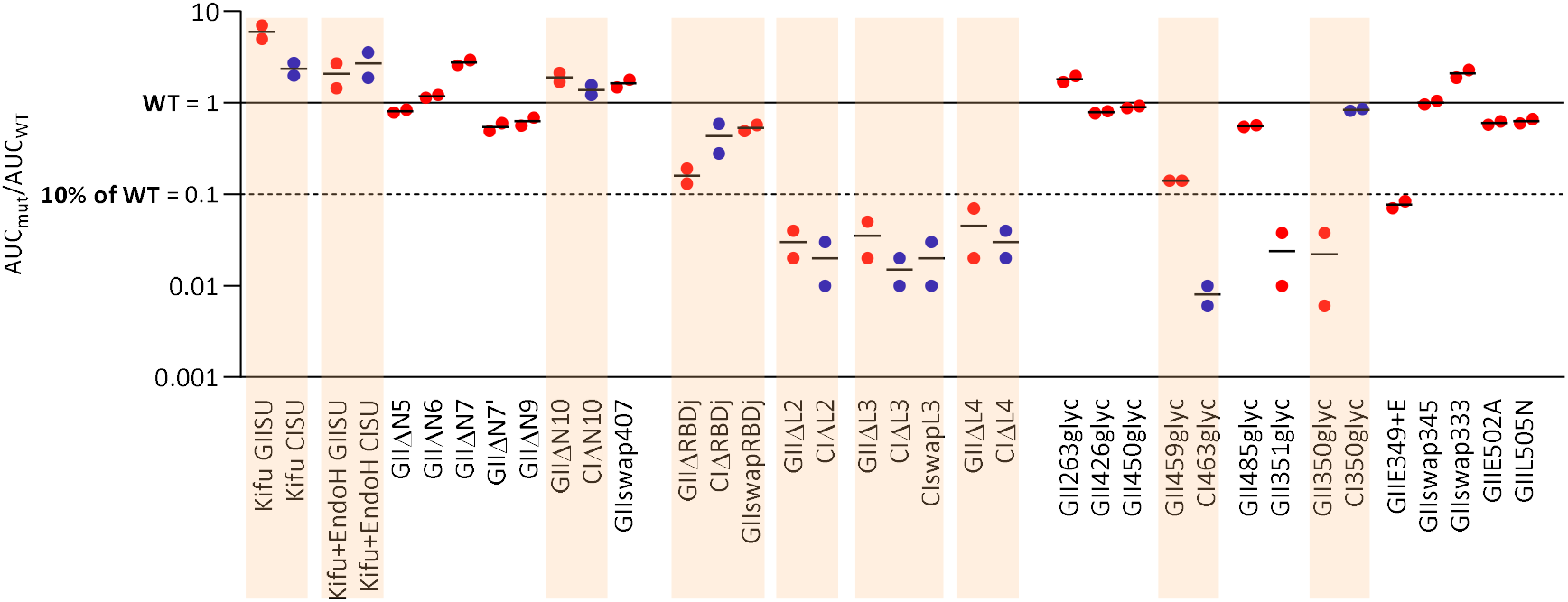
nAbs target epitopic regions involved in SU binding to susceptible cells or required for viral infectivity. A. HT1080 cells were incubated with the panel of tested immunadhesins (Fig. 3 to 7). Cell-bound SU were detected by staining with a fluorescently labeled antibody targeting the murine IgG Fc fragment. Stained cells were analyzed on a flow cell cytometer. Supplementary Fig. 8 presents the gating strategy and normalization of the results to the levels of bound WT SU (^GII^SU and ^CI^SU). Binding levels of GII mutants are presented by (red symbols). CI mutants (blue symbols) are presented side-by-side with the corresponding and related GII mutants, with a colored background. Mutated HBS shows reduced binding to susceptible cells, as shown in [14].

The binding data were also considered in the interpretation of nAb blocking experiments. The mutant SUs that retained their capacity to bind cells were considered to likely be properly folded and loss of nAb blockade could be attributed to the introduced mutation of the epitope. Overall, among all tested mutated immunoadhesins (Supplementary Table 1), only one failed to bind to cells and block the nAbs from all samples tested; thus the related data are not presented.

### Human plasma samples contain nAbs targeting a variable number of epitopic regions

Here, we show that nAbs recognize the three solvent-exposed loops on the upper RBD subdomain (Fig. 10D). We also identified mutations located C-terminal (^GII^459_glyc_ and ^CI^463_glyc_) of loop 4 that may affect nAb binding or SU folding. The glycosylation site N7’ had the greatest impact on the nAbs. It is located at the opening of the cavity in which the essential N8 glycan is buried [14]. Of note, the glycan inserted in ^GII^485_glyc_ may interfere with N8 function and/or accessibility to nAbs. In the RBD lower subdomain, we show the 345-353 loop to be an immunodominant epitope exclusively recognized by GII-specific nAbs and that it has a role in binding to cells. By contrast, the HBS was not frequently recognized.

**Fig. 10.**
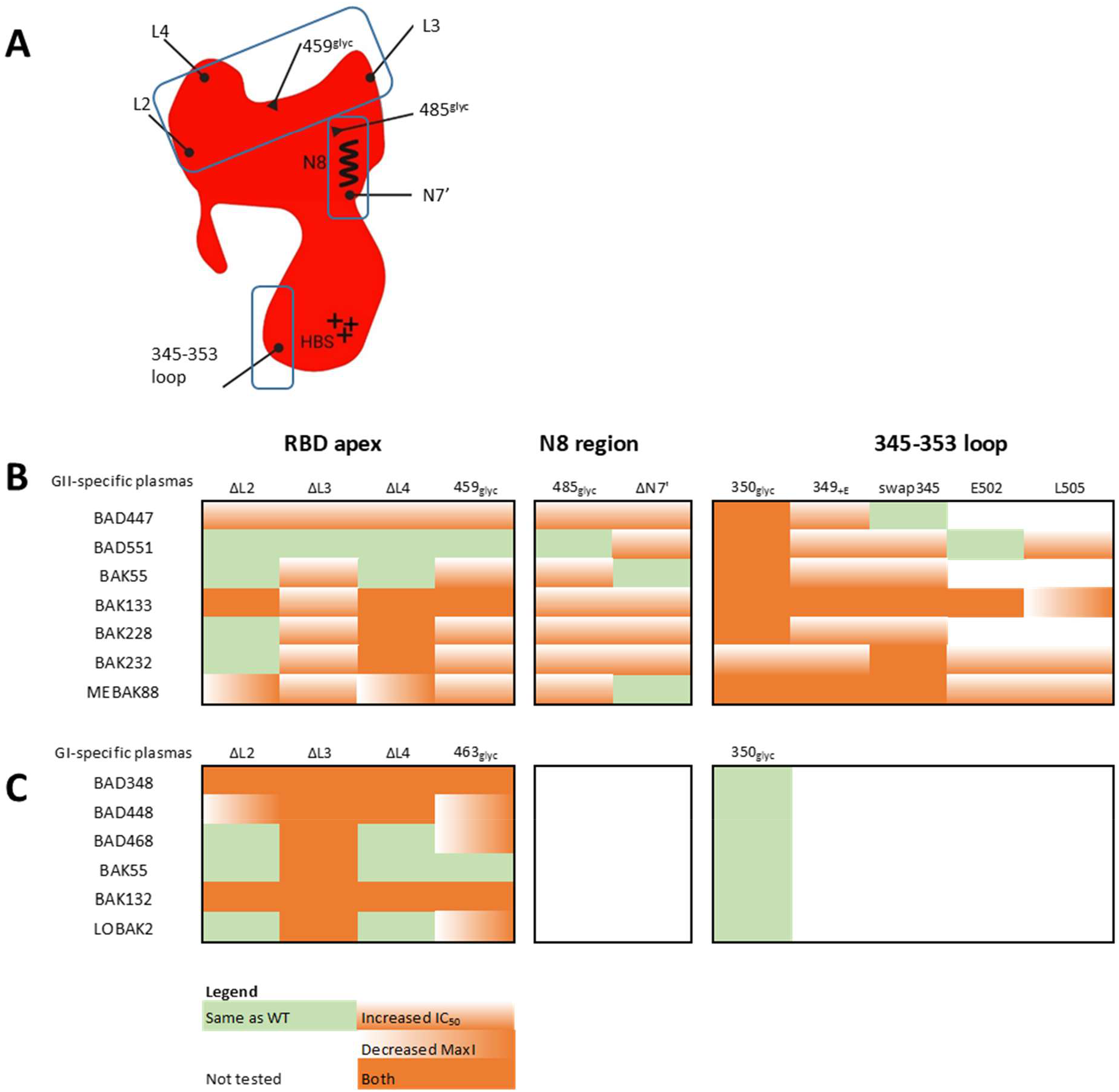
Human plasma samples contain nAbs targeting a variable number of epitopic regions. A. In the present study, we identified three epitopic sites on SUvar: the RBD apex, the N8 region, and the 345-353 loop. The schematic summary highlights L2, L3, and L4, the conserved N8 and adjacent N7’ glycosylation site, and the 345-353 loop (lines ending with a dot). Glycans inserted in predicted epitopes revealed additional antigenic sites (lines ending with a triangle). B and C. The results from competition experiments are summarized for each plasma sample and for the mutant SU from the most highly targeted regions (apex, N8 region, and the 345-353 loop). Four outcomes are presented: same recognition as WT SU (green), recognition with reduced affinity (i.e., increased IC_50_, orange vertical shading), blocking a smaller fraction of nAbs than WT SU (i.e., reduced MaxI, orange horizontal shading), or having both effects (orange). Black squares indicate the major epitopic regions of the GII- and GI-specific samples (panels B and C, respectively). Within each epitopic region defined by several mutant SU, nAb specificity varied between individuals.

The effect of the mutant SU on each plasma sample can be summarized as recognition similar to that of WT SU, recognition with a reduced affinity (i.e., increased IC_50_), blocking a smaller fraction of nAbs than WT SU (i.e., reduced MaxI), or having both effects. This summary highlights the two genotype-specific and immunodominant epitopic regions that we identified: loop 345-353 on ^GII^SU (Fig. 10B) and L3 on ^CI^SU (Fig. 10C). Within the RBD apex, four mutant SU correspond to different epitopic sites (L2, L3, L4, and 459/463^glyc^). GII-specific plasma samples presented interindividual variations in their specificity. As an example, one sample did not recognize the RBD apex (BAD551), whereas the six other samples did. In addition, samples differed in the number of targeted sites (BAK55 vs. BAK133), Similarly, GI-specific plasma samples contained nAbs focused on a single subdomain (such as BAK55 and ^CI^L3) or target epitopes on up to four sites (BAD348 and BAK132). We tested the recognition of the N8 region by GII-specific samples only and observed that all plasma samples were affected by alteration in the vicinity of N8. We examined the sequence of the identified epitopes and found that all were conserved on the SFV strains infecting individuals from this study or circulating in Central Africa (Supplementary Fig. 9). Overall, we identified three immunodominant epitopic regions (RBD apex, 345-353 loop, and N8 region), of which the sequences are conserved within each genotype. We provide evidence that SFV-infected individuals have nAbs that target several epitopes and recognize epitopes that differ between individuals.

## Discussion

Humans infected with zoonotic SFV raised potent neutralizing antibodies that target the genotype-specific SUvar domain overlapping most of the RDB. Here, we aimed to define the sites targeted by these antibodies. We demonstrate that nAbs raised by infected humans recognize three antigenic regions within the SUvar region from genotype II SFV. At least one of these regions is recognized by GI-specific nAbs. Based on the recently solved X-ray structure combined with functional data [13, 14], we hypothesize that the SFV-specific nAbs most likely target sites important for Env trimer formation and conformational changes that lead to membrane fusion, and/or sites important for binding to susceptible cells.

We identified only conformational epitopes, despite several attempts to capture linear epitopes using peptides (Fig. 8, Supplementary Fig. 7 and [24]). Our data are consistent with the low reactivity of NHP and human immune sera to denatured Env in immunoblot assays, which contain Abs that bind to native Env in radio-immunoprecipitation and neutralization assays [25-28]. Linear epitopes are usually located in mobile regions of the polypeptide chains. Such mobile segments are absent from the RBD lower subdomain formed by the compact assembly of α-helices and β-sheets [14]. In RBD monomers, the upper subdomain loops appear to be mobile, but inter-protomer interactions probably impose specific conformations in full-length Env. The loops likely form a number of discontinuous epitopes; L2 and L4 are in proximity in the monomer and L3 and L4 in the trimer [14].

Twelve plasma samples were used in this study, including one from an individual infected by strains belonging to both genotypes. Overall, the epitopic regions were recognized by all or most donors and can be considered to be immunodominant (Fig. 10). Importantly, within each genotype, sequences from the epitopic regions were identical among SFV strains infecting the studied individuals and those circulating in the same geographical area (Supplementary Fig. 9). This observation is consistent with the genetic stability of SFV [29]. The use of polyclonal plasma samples was key to providing a global picture of nAbs raised upon infection with zoonotic SFV, but only epitopes recognized by a significant fraction of plasma antibodies can be detected. Thus, future studies may identify subdominant epitopes and provide a more precise definition of those that are immunodominant.

SU proteins fully blocked nAbs from certain samples. However, for most tested samples, the infectivity was maintained but at levels lower than those observed for FVVs not exposed to plasma samples (Fig. 2). The heterogeneous glycosylation of the SU could explain the partial blockade of nAbs [27], as proposed for the incomplete neutralization of HIV [19-22]. The SU antigenicity of soluble monomeric proteins probably differed from those contained within viral particles due to the lack of the intra- and interprotomer interactions that form quaternary structures. Thus, our data identify nAb epitopes presented on the monomeric SU domain and indirectly suggest the existence of quaternary epitopes.

We have provided experimental evidence for the targeting of different epitopes by nAbs raised after infection by the two SFV genotypes. The RBD fold from different genotypes and FV from different host species is highly conserved [14]. In superinfection-resistance experiments, the CI Env inhibited entry from genotype II strains, indicating that SFV from different genotypes share the use of at least one molecule for entry into target cells; this molecule could act as an attachment factor or receptor [30]. The upper RBD subdomain was recognized by both GI- and GII-specific nAbs and the lower RBD subdomain by GII-specific nAbs only. GI-specific nAbs may, nevertheless, recognize epitopes on the lower RBD subdomain that are yet to be identified. Indeed, the biochemical properties of GI Env protein resulted in low expression and protein aggregation. Therefore, we used an SU from chimpanzee genotype I SFV to map epitopes recognized by antibodies induced by a gorilla genotype I SFV. We previously reported that plasma samples cross-neutralize both SFV species, with a strong correlation between nAb titers; however, nAb titers were globally higher against the GI than CI strain [7]. Thus, we may have missed a number of GI-specific epitopes.

Relative to other viruses, SFV-specific nAbs target a limited region on Env, i.e., they do not recognize epitopes on the TM nor non-RBD sites of the SU [31-33]. Based on the inhibition mechanisms described for other viruses [33], we can propose a number of possible modes of action. The targeting of RBD loops by nAbs may prevent the fusogenic conformational transition and exposure of the fusion peptide, which is located at the center of the trimer and shielded by the RBD, as visualized on viral particles [15]. It is possible that certain loop-specific nAbs bind several protomers, as described for HIV- and Ebola-specific nAbs [34, 35]. An alternative neutralization mechanism could be the targeting of epitopes located close to the protomer interface, resulting in trimer disruption, as described for HIV-1, SARS-CoV2 and Flu-specific nAbs [36-38].

Many nAbs interfere with particle binding to susceptible cells. The existence of a *bona fide* receptor for SFV is yet to be demonstrated. Consequently, the molecular determinants of Env binding to susceptible cells within the lower RBD subdomain are still only roughly defined. Certain nAbs recognize attachment factors, such as an HBS on SARS-CoV2 spike [39]. The SFV Env mutants deficient for HS binding efficiently blocked most plasma samples, indicating that HBS is either not a major nAb target or that the antigenicity of these mutants is preserved. Of note, the N7’ site located in the vicinity of HBS was recognized. All SU designed to map potential B-cell epitopes were tested for their capacity to bind susceptible cells. SU from the two genotypes differed in certain residues involved in cell binding. Most notably, nAbs recognizing the 345-353 loop may prevent GII Env binding, whereas the loop on GI Env is not involved in cell binding and is not targeted by nAbs (Fig. 9). This last observation highlights, for the first time, genotype-specific differences in SFV binding to susceptible cells and could be a starting point for further studies on identifying attachment factors and receptors for SFV.

Our choice of a functional strategy to map the epitopes targeted by nAbs proved to be critical due to the lack of linear epitopes. Overall, the use of human samples and soluble SU as competitor allowed us to create the first map of targeted regions. We considered several caveats (polyspecificity of plasma samples, absence of quaternary epitopes). We also carefully considered the possible nonspecific effect of mutations on protein folding, which could generate misleading results. Certain mutants were, indeed, poorly expressed and could not be used (Supplementary Table 1). A number of these limitations may be overcome in the future with the use of alternative tools, such as human monoclonal antibodies and subviral particles.

Human infection with zoonotic SFV represents a model for cross-species transmission of retroviruses leading to persistent infection that is successfully controlled by the immune system. We have previously reported the presence of potent nAbs in most infected individuals [24]. Here, we have mapped major antigenic sites. Concerning the control of SFV in humans, a notable result from the present study is that two immune escape mechanisms, sequence variation and glycan shielding, were not observed. The SFV RBD is structurally different from known and modelled retroviral RBDs [40]. We have provided a novel model integrating structural, genetic, functional, and immunological knowledge on the bimorphic SFV RBD. We have thus gained information on the two SFV genotypes that have persisted for over 30 million years of evolution with their animal hosts. Through the study of SFV and its unique properties, we should also gain fundamental knowledge on the structural basis for the inhibition of viruses by nAbs.

## Methods

### Human plasma samples

Blood samples were drawn from adult populations living in villages and settlements throughout the rainforests of Cameroon. Participants gave written informed consent. Ethics approval was obtained from the relevant national authorities in Cameroon (the Ministry of Health and the National Ethics Committee) and France (Commission Nationale de l’Informatique et des Libertés, Comité de Protection des Personnes Ile de France IV). The study was registered at www.clinicaltrials.gov, https://clinicaltrials.gov/ct2/show/NCT03225794/. SFV infection was diagnosed by a clearly positive Gag doublet on Western blots using plasma samples from the participants and the amplification of the integrase gene and/or LTR DNA fragments by PCR using cellular DNA isolated from blood buffy-coats [3]. We identified the SFV origin by phylogenetic analysis of the integrase gene sequence [3]. The SFV genotype was determined by amplification of the SUvar DNA fragment by PCR [7]. Plasma samples from 24 participants were used for this study (Supplementary Tables 2 and 4). Four participants were not infected with SFV and 20 were infected with a gorilla SFV.

### Viral strains, amino-acid numbering, and Env domain nomenclature

We used sequences from primary zoonotic gorilla SFVs, SFVggo_huBAD468 (GI-D468, JQ867465) and SFVggo_huBAK74 (GII-K74, JQ867464) [41] and the laboratory-adapted chimpanzee SFV, SFVpsc_huPFV (CI-PFV, KX087159) [42] for the synthesis of foamy viral vectors (FVV) and envelope proteins and peptides. For simple reference to previously described Env sequences and functions, we used the amino-acid positions from the CI-PFV strain, unless otherwise stated. The GI-D468, GII-K74, and CI-PFV sequence alignment are shown in supplementary Fig. 1. When referring to infecting SFV strains and the antibodies raised against them, gorilla and chimpanzee genotype I SFV are referred to as GI and CI, respectively; gorilla genotype II SFV is referred to as GII.

### Cells

Baby hamster kidney (BHK)-21 cells (ATCC-CLL-10, hamster kidney fibroblast) were cultured in DMEM-5% fetal bovine serum (FBS). HT1080 cells (ECACC 85111505, human fibrosarcoma) were cultured in Eagle’s Minimum Essential Medium with Earle’s Balanced Salts and L-glutamine supplemented with 10% FBS and 1% L-glutamine. Human embryonic kidney 293T cells (Cat. N° 12022001, Sigma) were cultured in DMEM-10% FBS. FreeStyle 293-F cells (Life Technologies) were cultured in Ex-cell 293 HEK serum-free medium supplemented with 5 µg/mL phenol red sodium salt, 2% L-glutamine, and 0.2x penicillin-streptomycin.

### Plasmids

The four-component CI-PFV FVV system (pcoPG, pcoPP, pcoPE, and puc2MD9) and the gorilla SFV Env constructs containing sequence from the zoonotic GI-D468 and GII-K74 *env* genes have already been described [7, 43]. Novel plasmids were synthesized by Genscript (Piscataway, NJ, USA). We built a FVV expressing β-galactosidase with a nuclear localization signal (puc2MD9-B-GAL) by replacement of the *gfp* gene in the puc2MD9 backbone for easier image analysis on our quantification device. GII-K74 Env with swapped 345-353 loop was constructed with the boundaries used for SU (Supplementary Table 1).

Immunoadhesin constructs express a fusion protein formed by the murine IgK signal peptide, SFV SU with deleted furin cleavage site (aa 127-567), and the heavy chain (hinge-CH2-CH3) of murine IgG2a. A Twin-Strep-tag was fused after the mIgG2a to facilitate immunoadhesin purification, except for the first immunoadhesin constructs (^GII^ΔRBDj, ^GII^swapRBDj and ^GII^ΔN10). The murine leukemia virus (MLV) gp70 SU (aa 34-475, strain FB29, NP_040334.1) was fused to the Twin-Strep-tag. All genes were codon optimized for expression in mammalian cells and placed under the control of a CMV promotor and intron in the pcZI plasmid [44].

The codon-optimized synthetic genes encoding GII-K74 SU (aa 127-566) and ectodomain (aa 91-907, [14]) were cloned into the already described pT350 plasmid [45], which contains the *Drosophila* metallothionein promoter, which is inducible by divalent cations [46], the *Drosophila* BiP signal peptide (MKLCILLAVVAFVGLSLG) at the N-terminus and a Twin-Strep-tag (AGWSHPQFEKGGGSGGGSGGGSWSHPQFEK) for affinity purification at the C-terminus. Stable S2 cell lines were generated by antibiotic resistance and protein production was induced by the addition of 2 µM CaCl_2_. For expression in mammalian cells, the murine IgK signal peptide was fused at the N-terminus of codon-optimized genes encoding GII-K74 SU (aa 127-566). The genes were placed under the control of the CMV promoter and intron [44] by insertion into the pcDNA3.1 vector. The ectodomain proteins with mutation in HBS are described in [14].

### Protein expression and purification

293-F cells were seeded at 2.5 10^6^ cells/mL in FreeStyle 293 Expression medium and transfected by the addition of plasmid DNA (2 µg/mL) and LipoD293 (6 µg/mL #SL100668, Tebu-bio). After 24 h, cells were diluted 1:1 in Ex-cell culture media and cultivated for another 48 h and the supernatants collected and stored at -80°C. For the expression of immunoadhesins with high-mannose-type glycans, transfected cells were cultivated in the presence of 5 µM Kifunensine mannosidase inhibitor (#BML-S114-0005, Enzo Life Sciences). Glycans were removed by incubating the immunoadhesins ON at room temperature (RT) with endoglycosidase H (endo-H, 0.8 U/mL, #P0702L, New England Biolabs) in 50 mM Na acetate-50 mM NaCl, pH 6. Supernatants were thawed and filtrated through a 0.2 µm filter before centrifugation at 4500 x g at 4°C in VivaSpin tubes (Sartorius). The first-generation immunoadhesins do not carry the Twin-Strep-tag and were purified using Protein A Mag Sepharose™ Xtra beads (50 µL beads/mL of supernatant; #28-9670-62, GE Healthcare) using binding buffer pH 7, 0.2 M Na_3_PO_4_ (#28-9030-59, Ab Buffer Kit, GE Healthcare). Beads were incubated at 4°C on a rotator ON. Proteins were eluted with a pH 2.5 elution buffer containing 0.2 M Glycine-HCl and neutralized with 1 M Tris-HCl, pH 9 (#PUR004, NEO Biotech, CliniSciencesProteus). Immunoadhesins and ^MLV^SU fused to Twin-Strep-tag were purified using Gravity Flow Strep-Tactin XT Superflow High-Capacity columns (#2-4031-001, Iba Lifescience). Concentrated proteins were resuspended in buffer W at pH 8 and incubated with BioLock (#2-0205-050, Iba Life sciences) for 15 min at RT to neutralize biotin. Loading onto the columns and elution were performed according to manufacturer’s instructions.

GII-K74 SU, WT, and mutated ectodomains were expressed in *Drosophila* Schneider’s cell line 2 (S2) cells following the standard protocol [47]. GII monomeric SU and trimeric ectodomain proteins were purified by affinity chromatography on a Strep-Tactin column (IBA Biosciences), followed by size-exclusion chromatography on Superdex 200, using standard methods provided by the manufacturers.

Protein concentrations were measured by NanoDrop. To verify protein purity and aggregate formation, 1.5 µg of purified proteins was heat-denaturated at 70°C for 10 min in NuPAGE LDS sample buffer (#NP0007, Introgen), with or without NuPAGE reducing buffer (50 mM DTT; #NP0009, Invitrogen). Samples were loaded onto a precast NuPAGE 4-12% Bis-Tris gel (#NP0322, Invitrogen). The gel was incubated with Bio-Safe Coomassie staining solution (#1610786, Bio-Rad) for 2 h with gentle shaking, rinsed in H_2_O ON, and imaged using a G:BOX (Syngene) (Supplementary Fig. 4).

### Western blots

Western-blot analysis of protein expression was performed on cell supernatants. Samples were heat-denaturated and reduced as described for purified proteins before loading onto precast NuPAGE 4-12% Bis-Tris gels (#NP0322, Invitrogen). Proteins were migrated for 2-3 h at 120 V in NuPAGE MOPS SDS running buffer (#NP0001, Invitrogen) and transferred onto a PVDF membrane (#1704156, Bio-Rad) using a Trans-Blot Turbo transfer system (Bio-Rad). Membranes were blocked and antibody labeled in Tris-buffered saline (TBS)-0.1%Tween-5% BSA. For Strep-tag detection, membranes were incubated with StrepMAB-Classic monoclonal antibody conjugated to HRP (0.05 µg/mL, #2-1509-001, Iba Lifesciences) for 2 h at RT. For SFV SU detection, membranes were incubated with a biotinylated anti-SU murine antibody ON at 4°C (3E10, 1 µg/mL), washed three times in TBS-0.1% Tween for 10 min and incubated with Streptavidin-HRP (1:2000, #DY998, Biotechne). Membranes were washed three times in TBS-0.1% Tween for 10 min before revelation with SuperSignal West Pico PLUS chemiluminescence substrate (#34579, ThermoFischer Scientific) and signal acquisition using a G:BOX.

### Foamy viral vectors (FVVs)

FVV particles were produced by co-transfection of HEK293T cells by the four plasmids. Fifteen µg total DNA (gag:env:pol:transgene ratio of 8:2:3:32) and 45 µl polyethyleneimine (#101-10N, JetPEI, Polyplus) were added to a 10 cm^2^ culture dish seeded with 4 × 10^6^ 293T cells. Supernatants were collected 48 h post-transfection, clarified at 500 x g for 10 min, filtered using a 0.45 µm filter, and stored as single-use aliquots at -80°C. FVVs were titrated as described [7, 48], with minor modifications to optimize X-Gal staining of transduced cells, which was lighter than that of infected cells. Briefly, FVV samples were thawed and clarified by spinning at 10,000 x g at 4°C for 10 min. Serial five-fold dilutions were added in triplicate to sub-confluent BHK-21 cells seeded in 96-well plates and cultured for 72 h at 37°C. Plates were fixed with 0.5% glutaraldehyde in PBS for 10 min at RT, washed with PBS, and stained with 150 µl X-gal solution containing 2 mM MgCl_2_, 10 mM potassium ferricyanide, 10 mM potassium ferrocyanide, and 0.8 mg/mL 5-bromo-4-chloro-3-indolyl-B-D-galactopyranoside in PBS for 3 h at 37°C. Blue-stained cells were counted using an S6 Ultimate Image UV analyzer (CTL Europe, Bonn, Germany). One infectious unit was defined as one blue cell. Cell transduction by FVV is a surrogate for viral infectivity and FVV titers are expressed as infectious units (IU)/mL.

The yield of FVV particles was estimated by quantifying particle-associated transgene RNA. FVV RNA was extracted from raw cell supernatants using a QIAamp Viral RNA Extraction Kit (Qiagen), treated with a DNA-free kit (Life Technologies), and retro-transcribed with Maxima H Minus Reverse Transcriptase (Thermo Fischer Scientific) using random primers (Thermo Fischer Scientific) according to the manufacturers’ instructions. qPCR was performed on cDNA using BGAL primers (BGAL_F 5’ AAACTCGCAAGCCGACTGAT 3’ and BGAL_R 5’ ATATCGCGGCTCAGTTCGAG 3’) with a 10-min denaturation step at 95°C and 40 amplification cycles (15 s at 95°C, 20 s at 60°C, and 30 s at 72°C) carried out using an Eppendorf realplex2 Mastercycler (Eppendorf). A standard curve prepared with serial dilutions of puc2MD9-BGAL plasmid was used to determine the FVV copy number. Results are expressed as vector particles/mL, considering that each particle carries two copies of the transgene.

To test the capacity of mutated Env to mediate the binding of vector particles to cells, HT1080 cells were incubated with the FVV particles (1, 10 and 100 particles/cell) on ice for 1 h. The cells were washed three times with PBS to eliminate unbound FVV and RNA was extracted using an RNeasy plus mini-Kit (Qiagen) according to manufacturer’s protocol and RT performed as described for FVV RNA quantification. Bound FVV was quantified by qPCR of the *bgal* gene as described for vector titration; cells were quantified by qPCR amplification of the *hgapdh* gene with the following primers: hGAPDH_F 5’ GGAGCGAGATCCCTCCAAAAT 3’ and hGAPDH_R 5’ GGCTGTTGTCATACTTCTCATGG 3’. The qPCR reaction conditions were the same as those used to amplify the *bgal* gene. The relative mRNA expression of *bgal* versus *hgapdh* was calculated using the -ΔΔCt method, and relative binding as 2-ΔΔCt.

### Neutralization assays

Prior to use in neutralization assays, plasma samples were diluted 1:10 in DMEM + 1 mM MgCl_2_, decomplemented by heating at 56°C for 30 min, and frozen at -80°C as single-use aliquots. Thawed plasma samples were clarified by spinning at 10,000 x g for 10 min at 4°C. Serial two-fold dilutions were incubated with FVV for 1 h at 37°C before titration in triplicate and residual infectivity quantified using the 96-well plate microtitration assay described above. P96-well plates seeded with 5,000 BHK-21 cells were exposed to 300 IUs of FVV. Neutralization titers were defined as the inverse of the plasma dilution required to reduce viral infectivity by half. Plasma samples were initially titrated against replicating virus [7]. We selected those with neutralization titers > 1:100 against the virus and performed plasma titration against the FVV. We defined the plasma dilution required for a 90% reduction of FVV infectivity and used it as the fixed concentration for the nAbs. Plasma samples were incubated with serial dilutions of recombinant Env proteins for 1 h at 37°C. FVV was then mixed with the plasma/protein preparation and incubated 1 h at 37°C before addition to BHK-21 cells. FVV infectivity was quantified 72 h later as described for their titration. All conditions were tested in triplicate and the mean IU/well calculated. Cells transduced with mock treated FVV (i.e., incubated with ^MLV^SU at 20 nM or with medium) were quantified on each plate and this value used as a reference. Relative infectivity was calculated for wells treated with the plasma/protein mix and is expressed as the percentage of the reference value. The WT SU (^GII^SU or ^CI^SU) matched with the FVV Env was titrated along with the tested mutant SU on every plate in every experiment for each plasma tested. All SU were tested against each plasma at least twice and against at least four plasma samples. In the first assay (screening), SU were added at three concentrations, ranging from 200 to 2 nM. In the second assay (confirmation), SU with activity similar to that of WT or with no activity were tested a second time at three dilutions. Mutant SU with intermediate activity were titrated in a three-fold serial dilution setting starting at 60 nM. We used two parameters to define the action of the SU on the nAbs from each plasma sample: maximum infectivity (MaxI) and 50% inhibitory concentration (IC_50_). MaxI corresponds to the maximal infectivity in the presence of SU-blocking nAbs and is defined as the mean relative infectivity at the two highest doses tested. The IC_50_ values were calculated by plotting the relative infectivity as a function of SU concentration and a four parameters regression modeling of the Prism software (Version 9, GraphPad). Mutant SU with nAb activity similar to the WT one were given its IC_50_, and those with minimal or no activity were given an arbitrary IC_50_ value of 200 nM, corresponding to the highest concentration of SU tested.

### SFV Env binding to cells

SU were thawed at RT and clarified at 10,000 x g for 10 min. HT1080 cells were treated with trypsin-EDTA and 5 × 10^5^ cells resuspended in 0.1 mL FACS buffer (PBS-0.1% BSA) containing the SU and incubated on ice for 1 h. Cells were washed twice and incubated with donkey anti-murine IgG-(H+L)-AF488 (20 µg/mL, #A21202, Invitrogen) on ice for 30 min. Cells were washed and resuspended in 0.2 mL PBS-2% PFA. Data were acquired on a CytoFlex cytometer (Beckman Coulter) and analyzed using Kaluza software. A minimum of 25,000 cells were acquired. Single cells were selected by sequential gating on FSC-A/SSC-A and SSC-A/SSC-H dot-plots (Supplementary Fig. 8). SU binding is expressed as the ratio of mean fluorescence intensity (MFI) of SU-treated over untreated cells. Each SU was tested twice at 3, 30, and 300 nM. The MFI ratios were plotted as a function of SU concentration and the area under the curve was calculated (Supplementary Fig. 8). To limit inter-experimental variation, WT SU were included in every experiment and the binding level of mutant SU normalized to that of the WT.

### Peptides

The SUvar (aa 248-488) sequences from the GI-D468 and GII-K74 strains were analyzed using linear B-cell epitope prediction software available on the Immune Epitope Data Base (http://tools.iedb.org/bcell/), which are based on known B-cell epitopes (LBtope [49]), hydrophilicity (Parker prediction replaced by Bepipred software [50]), and protrusion outside of globular proteins (Ellipro [51]). In addition, we manually inspected viral sequences for stretches of genotype-specific epitopes. We selected 14 sequences and tested the corresponding GI-D468 and GII-K74 peptides, as well as nine CI-PFV peptides from a previously synthesized peptide set [24] (Supplementary Table 5). After resolution of the RBD crystal structure, we designed a second peptide set corresponding to the four loops located in the RBD upper subdomain [14]. As positive controls, we used the N_96_-V_110_ (NKDIQVLGPVIDWNV from SFV Env LP) and I_176_-I_199_ (INTEPSQLPPTAPPLLPHSNLDHI from HTLV-1 gp46) peptides containing immunodominant epitopes [24]. Peptides were synthesized by Smartox SAS (Saint-Martin d’Hères, France) or Genscript (Leiden, The Netherlands) and were tested individually.

### ELISA

The protocol was described in [24]. Briefly, peptides diluted in carbonate buffer at 1 µg/mL were coated on clear high-binding polystyrene 96-well microplates (Biotechne) overnight (ON) at +4°C. Plasma samples were diluted at 1:200 in phosphate buffered saline (PBS)-0.1% bovine serum albumin (BSA)-0.1% Tween20. Bound plasma antibodies were detected with horseradish peroxidase (HRP)-conjugated goat anti-human IgG H+L (0.02 µg/mL, #109-035-008, Jackson Immuno Research Europe). The peptide diluent was used as the negative control and antibody binding to peptides is expressed as the difference in OD (Δ_OD_ = OD_test_ – OD_control_). Plasma samples from three SFV-uninfected individuals were tested for binding to the 37 peptides. The mean + 2 SD of Δ_OD_ (0.14) was used to define the positivity cutoff. The proportion of responding individuals was calculated among those infected with a given virus (SFV, n = 17; HTLV-1, n = 7).

### Statistical analysis

Mutations in the SU modified their ability to block the nAbs in two ways: either by reducing avidity, leading to a higher IC_50_, or by the fraction of plasma nAbs blocked, leading to lower maximal infectivity. For each plasma sample, the IC_50_ and MaxI were tested for the WT SU in at least five experiments. The threshold value defining a statistically significant change relative to WT was defined as the mean + 3*SD for the IC_50_ and the mean -3*SD for MaxI. Thresholds were defined for each plasma sample. The infectious titers, particle concentration, percentage of infectious particles, and quantity of bound FVV carrying WT and mutant SU were compared using the paired t test (GraphPad Prism 9 software).

## Acknowledgements

We thank H. Mouquet and V. Lorin for sharing the 293-F cells and advice. We are grateful to Mathilde Couteaudier for mentoring MD and LTD in their performance of the experimental work. We used devices from the Cytometry and Biomarker Utechs at the Institut Pasteur and we thank their staff for their support and advice. We thank Delphine Brun from the Unité de Virologie Structurale for technical assistance. The manuscript has been edited by a native English speaker.

## Authors contribution

FB, MB, FR, and AG designed the project. FB and MB supervised the project and acquired funding. LTD, IF, YC, MD, and TM performed the experiments. LTD, IF, FB, and MB analyzed the data. AG, RN, and CBN collected the human samples. LTD and YC provided figures and text elements for the draft. FB wrote the initial and final drafts. All authors participated in reviewing the drafts.

