## Supplementary information for "Neutralization of zoonotic retroviruses by human antibodies: genotype-specific epitopes within the receptor-binding domain from simian foamy virus"

### *Dynesen et al.* Supplementary data

#### Supplementary Table 1. SFV Env proteins produced for the study

| Name | Description[a] | Expression level[b] | Coomassie gel shown in Supplementary Fig. 4 | Neutralization experiments presented in | Comment |
|---|---|---|---|---|---|
| **Immunoadhesins** | (SU fused to murine Fc and Strep-tag | | | | |
| $^{CI}$SU | WT immunoadhesin, CI-PFV strain | | Panel A | Fig. 7 | |
| Kifu $^{CI}$SU | WT immunoadhesin produced in the presence of Kifunensine | Normal | Panel B | Fig. 7 | |
| Kifu+Endo-H $^{CI}$SU | WT immunoadhesins produced in the presence of Kifunensine and treated with Endo-H | Normal | Panel B | Fig. 7 | |
| $^{CI}$ΔN10 | N423>A | Normal | Panel E | Fig. 7 | |

| | | | | |
|---|---|---|---|---|
| $^{CI}$ΔRBDj | ΔF397-S483 | Normal | Panel E | Fig. 7 |
| $^{CI}$ΔL2 | ΔK278-Y293 | Reduced | Panel F | Fig. 7 |
| $^{CI}$ΔL3 | ΔI411-R436 | Normal | Panel F | Fig. 7 |
| $^{CI}$swapL3 | CI-I411-R436>GII-V410-R433 | Normal | Panel F | Fig. 7 |
| $^{CI}$ΔL4 | ΔE445-P461 | Normal | Panel F | Fig. 7 |
| $^{CI}350_{glyc}$ | G350N | Normal | Panel F | Fig. 7 |
| $^{CI}352_{glyc}$ | S352N | Undetectable | | Fig. 7 |
| $^{CI}463_{glyc}$ | W463N | Normal | Panel F | Fig. 7 |
| $^{GII}$SU | WT immunoadhesin, GII-K74 strain | | Panel A | Fig. 3, 4, 5 |
| Kifu $^{GII}$SU | WT immunoadhesin produced in the presence of Kifunensine | Normal | Panel B | Fig. 3 |
| Kifu+Endo-H $^{GII}$SU | WT immunoadhesin produced in the presence of Kifunensine and treated with Endo-H | Normal | Panel B | Fig. 3 |
| $^{GII}$ΔN5 | N286>A | Reduced | Panel C | Fig. 3 |

| | | | | | |
|---|---|---|---|---|---|
| $^{GII}$ΔN6 | N311>A | Normal | Panel C | Fig. 3 | |
| $^{GII}$ΔN7 | N346>A | Normal | Panel C | Fig. 3 | |
| $^{GII}$ΔN7’ | N373>A | Reduced | Panel C | Fig. 3 | |
| $^{GII}$ΔN9 | N404>A | Reduced | Panel C | Fig. 3 | |
| $^{GII}$ΔN10 | N411>A | Reduced | Panel C | Fig. 3 | |
| $^{GII}$ΔN9N10 | N404>A + N411>A | Insufficient | Panel C | - | Aggregates |
| $^{GII}$swap407 | 407RYNVNET413>KDIKSEI | Reduced | Panel D | Fig. 3 | |
| $^{GII}$ΔRBDj | ΔF396-G480 | Normal | Panel E | Fig. 4 | |
| $^{GII}$swapRBDj | GII-F396-G480>GI-F396-A482 | Reduced | Panel E | Fig. 7 | |
| $^{GII}$ΔL2 | ΔK278-Y293 | Reduced | Panel F | Fig. 4 | |
| $^{GII}$ΔL3 | ΔV410-R433 | Reduced | Panel F | Fig. 4 | |
| $^{GII}$ΔL4 | ΔE442-P458 | Normal | Panel F | Fig. 4 | |
| $^{GII}263^{glyc}$ | D263N | Normal | Panel G | Fig. 4 | |
| $^{GII}426^{glyc}$ | H428>T | Normal | Panel I | Fig. 4 | |
| $^{GII}450^{glyc}$ | D450>N | Normal | Panel I | Fig. 4 | |
| $^{GII}$A438-A443 | 438-REGKKE-443>AAGAAA | Insufficient | | | |

| | | | | | |
|---|---|---|---|---|---|
| $^{GII}459^{glyc}$ | E459>N | Normal | Panel I | Fig. 4 | |
| $^{GII}485^{glyc}$ | E485>N | Normal | Panel I | Fig. | |
| $^{GII}364^{glyc}$ | K364>N + G366>T | Reduced | Panel I | | No nAb blocking and no cell binding |
| $^{GII}351^{glyc}$ | L353T | Reduced | Panel H | Fig. 5, Supplementary Fig. 5 | Aggregates; experiments repeated with SEC purified immunoadhesin |
| $^{GII}350^{glyc}$ | G350>N + K352>S | Reduced | Panel I | Fig. 5 | Aggregates, moderate |
| $^{GII}349_{+E}$ | E inserted afterT348 | Reduced | Panel I | Fig. 5 | |
| $^{GII}$swap333 | GII-L333-S345>CI-EQNERFLLNKLN | Reduced | Panel D | Fig. 5, Supplementary Fig. 6 | |
| $^{GII}$swap345 | GII-S345-N351>GI-NNLTELTS | Normal | Panel D | Fig. 5, Supplementary Fig. 6 | |
| $^{GII}$ΔT348-L353 | ΔT348-L353 + GG | Undetectable | | | |
| $^{GII}$swap349 | GII-I349-N355>CI-SGTSVLKK | Insufficient | | | |

| | | | | | |
|---|---|---|---|---|---|
| $^{GII}$E502A | E502>A | Reduced | Panel D | Fig. 5 | |
| $^{GII}$L505N | L505>N | Reduced | Panel D | Fig. 5 | |
| **Tagged proteins** | Proteins fused to a Strep-tag | | | | |
| $^{MLV}$SU | MLV SU, strain FB29 | Not applicable | A | Fig. 2 | Three bands are visible in non-reduced conditions that correspond to oligomers formed by the free cysteine thiol group [52]. |
| GII-K74 SU(m) | WT SU produced in mammalian cells | Not applicable | nd | Supplementary Fig. 2 | Purified by size exclusion chromatography [14] |
| GII-K74 SU (i) | WT SU produced in insect cells | Not applicable | nd | Supplementary Fig. 2 | Purified by size exclusion chromatography [14] |
| GII-K74 Ecto(i) | Env 91-907, produced in insect cells | Not applicable | nd | Supplementary Fig. 2 | Purified by size exclusion chromatography [14] |
| $^{GII}$K342A/R343A | K342>A+R343>A into GII-Ecto(i) | Normal | nd | Fig. 5 | Purified by size exclusion chromatography [14] |

| | | | | | |
|---|---|---|---|---|---|
| [GII]R356A/R369A | R356>A+R369>A into GII-Ecto(i) | Normal | nd | Fig. 5 | Purified by size exclusion chromatography [14] |

[a] aa positions indicated are those of each protein; for GII-K74, some differ from the CI-PFV based numbering (Supplementary Fig. 1). The symbols > and Δ designate aa substitutions and deletions, respectively.

[b] The level of expression was assessed on crude supernatants of transfected cells and categorized as undetected, insufficient to perform the experiments, decreased, or normal relative to the WT counterpart.

#### Supplementary Table 2. Plasma samples used for the neutralization study

| Participant | Ethnicity | SFV infection[a] |
|---|---|---|
| BAD448 | Bantu | GI |
| BAK132 | Pygmy | GI |
| LOBAK2 | Pygmy | GI |
| BAD551 | Bantu | GII |
| BAK133 | Pygmy | GII |
| BAK228 | Pygmy | GII |
| BAK232 | Pygmy | GII |
| MEBAK88 | Pygmy | GII |
| BAD348 | Bantu | GI+GII |
| BAD447 | Bantu | GI+GII |
| BAD468 | Bantu | GI+GII |
| BAK55 | Pygmy | GI+GII |

[a] Participants were infected with a gorilla SFV of which the genotype was defined by PCR using primers located within SUvar [7]. Among the four individuals infected by both genotypes, only one (BAK55) was tested against both genotypes in epitope mapping experiments because his nAb titers were high against both genotypes; the three other samples were tested against a single viral genotype.

#### Supplementary Table 3: Methods used to predict epitopic regions and to design the mutant SU proteins

| Name | Prediction | Genotype-specific features |
|---|---|---|
| $^{GII}$ΔN5, $^{GII}$ΔN6, $^{GII}$ΔN7, $^{GII}$ΔN9 | Functional study [23] | |
| $^{GII}$ΔN7' | Presence varied according to the viral strain [11] | Absent from CI-PFV strain |
| $^{GII}$ΔN10, $^{CI}$ΔN10 | Functional study [23] | Genotype-specific localization of N10 [11] |
| $^{GII}$swap407 | Genotype-specific sequence before N10 | Genotype-specific localization of N10 [11] |
| $^{GII}$ΔRBDj, $^{GII}$swapRBDj, $^{CI}$ΔRBDj, $^{CI}$swapRBDj | Functional study [13] | $^{GII}$SU with $^{CI}$RBDj ($^{GII}$swapRBDj) was expressed; $^{CI}$SU with $^{GII}$RBDj was not |
| $^{GII}$K342A/R343A, $^{GII}$R356A/R369A | Functional study [14] | Mutations introduced in GII-K74 ectodomain; CI-PFV ectodomain is expressed at low levels |
| $^{GII}$ΔL2, $^{CI}$ΔL2 | Crystal structure [14] + disordered secondary structure[a] | |
| $^{GII}$ΔL3, $^{CI}$ΔL3, $^{CI}$L3swap, $^{GII}$ΔL4, $^{CI}$ΔL4 | Crystal structure ([14], cosubmitted) | |
| $^{GII}263^{glyc}$, $^{GII}426^{glyc}$, $^{GII}450^{glyc}$, $^{GII}459^{glyc}$ | Genotype specific + CBtope[b] | |
| $^{GII}351^{glyc}$, $^{GII}364^{glyc}$, $^{GII}485^{glyc}$ | Genotype specific + CBtope[b] + disordered secondary structure[a] | Loop size around aa351 differs between genotypes |

| | |
|---|---|
| $^{GII}350^{glyc}$, $^{GII}349_{+E}$, $^{GII}$swap345, $^{GII}$swap333, $^{GII}$E502A, $^{GII}$L505N, $^{CI}$G350$^{glyc}$ | Designed after testing $^{GII}351^{glyc}$ |
| $^{CI}463^{glyc}$ | Designed after testing GII $^{GII}459^{glyc}$ |

[a] The Protein Homology/analogY Recognition Engine V 2.0 (Phyre2) web portal was used for secondary structure prediction [53]. Disordered secondary structures predicted by the software were considered to define epitopic regions to be tested.

[b] CBtope software predicts conformational B-cell epitopes on the basis of their primary sequence (http://crdd.osdd.net/raghava/cbtope/, [54]). We applied the recommended parameters, i.e. 19 aa-long window, -0.3 threshold, and considered values ≥ 4 as potential epitopes.

#### Supplementary Table 4. Plasma samples used for the ELISA assays

| Participant | Ethnicity | SFV infection[a] | Fig. 8[b] | Supplementary Fig. 7[b] |
|---|---|---|---|---|
| BAD356 | Bantu | Uninfected | | X |
| BAK141 | Pygmy | Uninfected | | X |
| BAK183 | Pygmy | Uninfected | | X |
| BAK279 | Pygmy | Uninfected | | X |
| MEBAK195 | Pygmy | GI | X | |
| BAD448 | Bantu | GI | X | X |
| BAD463 | Bantu | GI | X | |
| BAK132 | Pygmy | GI | X | X |
| BAK56 | Pygmy | GI | X | |
| BAK82 | Pygmy | GI | X | |
| LOBAK2 | Pygmy | GI | X | X |
| BAD551 | Bantu | GII | X | X |
| BAK133 | Pygmy | GII | | X |
| BAK232 | Pygmy | GII | X | X |
| MEBAK88 | Pygmy | GII | | X |
| BAD348 | Bantu | GI+GII | X | |
| BAD447 | Bantu | GI+GII | X | |
| BAD468 | Bantu | GI+GII | X | X |
| BOBAK153 | Pygmy | GI+GII | X | |
| BAD456 | Bantu | GI+GII | X | |
| BAK177 | Pygmy | GI+GII | X | |
| BAK55 | Pygmy | GI+GII | X | |
| BAK74 | Pygmy | GI+GII | X | |

[a] Participants were infected with a gorilla SFV of which the genotype (GI or GII) was defined by PCR using primers located within SUvar [7]. Some participants were coinfected by strains from both genotypes (GI+GII). [b] The samples used are indicated for each of the two sets of peptides tested and presented in Fig.8 and supplementary Fig. 7.

#### Supplementary Table 5. Synthetic peptides used to search for linear epitopes

| Peptide name | Position[a] | Sequence[b] | Length | Prediction[c] |
|---|---|---|---|---|
| BAD468-247 | 247-270 | $_{RR}$PSEELIADQCPLPGYHAGVEYTTQ$_{RR}$ | 28 | Parker |
| BAD468-267 | 267-284 | RYTTQAIWDYYIKVEITRP | 19 | Genotype-specific |
| BAD468-280 | 280-300 | EITRPKNWTSYAQYGNARLGS$_{R}$ | 22 | Parker + Lbtope |
| BAD468-308 | 308-337 | RKNFTHVLFCSDQLYAKWYNIENTLLKNEE$_{R}$ | 31 | Ellipro |
| BAD468-329 | 329-342 | $_{R}$ENTLLKNEELLQKK | 15 | Parker |
| Bad468-340 | 340-357 | LQKKLNNLTELTSLLKKR | 18 | Genotype-specific |
| BAD468-350 | 350-374 | TSLLKKRALPRTWTTQGKNNLFRNI | 25 | Lbtope |
| BAD468-399 | 399-418 | RWEGDCNYTKDKISEIVPQCKR | 22 | Parker |
| BAD468-411 | 411-432 | RSEIVPQCKGFYNNSKWMHMHPYR | 24 | Parker |
| BAD468-425 | 424-444 | SKWMHMHPYACRFWRNKNEKE | 21 | Lbtope |
| BAD468-435 | 435-454 | RFWRNKNEKEETKCDGRDDN | 20 | Lbtope |
| BAD468-441 | 441-460 | RNEKEETKCDGRDDNKCLYYPRR | 23 | Parker + Lbtope |
| BAD468-450 | 450-472 | RRGRDDNKCLYYPLWDSPEATYDFGRRR | 28 | Parker + Lbtope |
| BAD468-487 | 487-512 | SSKQIRQQDYEVYSIYQECKLASRIH | 26 | Lbtope |
| BAK74-247 | 247-270 | RRPNEGLIADQCPLPGLADVSFYPYQRRR | 29 | Parker |
| BAK74-267 | 267-284 | RYPYQAIWDYYAKIENIRP | 19 | Genotype-specific |
| BAK74-280 | 280-300 | RENIRPANWTSSKLYGKARMGSR | 23 | Parker + Lbtope |
| BAK74-308 | 308-338 | RNINNTHILFCSDVLYSKWYNLQNSILQNENR | 32 | Ellipro |
| BAK74-330 | 330-348 | RQNSILQNENELTKRLSNLT | 20 | Parker |
| Bak74-339 | 339-355 | ELTKRLSNLTIGNKLKN | 17 | Genotype-specific |
| BAK74-350 | 350-374 | GNKLKNRALPYEWAKGGLNRLFRNI | 25 | Lbtope |
| BAK74-399 | 399-418 | RWEGDCNITRYNVNETVPECKR | 22 | Parker |
| BAK74-411 | 411-430 | RNETVPECKDFPHRRFNDHPYR | 22 | Lbtope |
| BAK74-425 | 424-442 | RRFNDHPYSCRLWRYREGKE | 20 | Lbtope |
| BAK74-435 | 433-452 | RLWRYREGKEEVKCLTSDHTR | 21 | Lbtope |
| BAK74-441 | 439-458 | REGKEEVKCLTSDHTRCLYYPRR | 23 | Parker + Lbtope |
| BAK74-449 | 449-470 | RRSDHTRCLYYPEYSNPEALFDFGRR | 26 | Parker + Lbtope |
| BAK74-485 | 485-510 | RESTSIRQQDYEVYSIYQECKLASKTYR | 28 | Lbtope |
| PFV-37 | 251-265 | LIADQCPLPGYHAGL | 15 | Genotype-specific |
| PFV-41 | 271-285 | SIWDYYIKVESIRPA | 15 | Genotype-specific |
| PFV-46 | 296-310 | ARLGSFYIPSSLRQI | 15 | Genotype-specific |
| PFV-55 | 341-355 | LNKLNNLTSGTSVLK | 15 | Genotype-specific |
| PFV-65 | 391-405 | NTSYYSFSLWEGDCN | 15 | Genotype-specific |
| PFV-66 | 396-410 | SFSLWEGDCNFTKDM | 15 | Genotype-specific |
| PFV-80 | 466-470 | PESTYDFGYLAYQKN | 15 | Genotype-specific |
| PFV-81 | 471-485 | DFGYLAYQKNFPSPI | 15 | Genotype-specific |
| PFV-82 | 476-490 | AYQKNFPSPICIEQQ | 15 | Genotype-specific |
| L1-GI | 258-269 | $_{R}$LPGYHAGVEYTT$_{R}$ | 14 | RBD structure |
| L1-GII | 258-269 | $_{R}$LPGLADVSFYPY$_{R}$ | 14 | RBD structure |
| L2-GI | 279-288 | VEITRPKNWT$_{R}$ | 11 | RBD structure |
| L2-GII | 279-288 | IENIRPANWT$_{R}$ | 11 | RBD structure |
| L3-GI | 411-435 | SEIVPQCKGFYNNSKWMHMHPYACR | 25 | RBD structure |
| L3-GII | 410-433 | VNETVPECKDFPHRRFNDHPYSCR | 24 | RBD structure |
| L4-GI | 447-457 | $_{R}$KCDGRDDNKCL | 12 | RBD structure |
| L4-GII | 445-455 | $_{R}$KCLTSDHTRCL | 12 | RBD structure |

[a] Positions refer to each viral sequence.
[b] Subscript characters indicate residues added to increase peptide solubility.
[c] Linear B-cell epitopes were predicted using the software available on the Immune Epitope Data Base (http://tools.iedb.org/bcell/): LBtope ([49] and Parker hydrophilicity prediction replaced by the Bepipred program [50] and Ellipro [51]). Genotype-specific sequences were manually defined. After resolution of the RBD structure [14], eight novel peptides overlapping the four loops were synthesized.

### Supplementary Fig. 1. CI-PFV, GI-D468, and GII-K74 Env sequence alignment

```
               |LP
CI-PFV         MAPPMTLQQW IIWKKMNKAH EALQNTTTVT EQQKEQIILD IQNEEVQPTR    50
GI-D468        .....S.... ...N..HQ.. Q....S.L.. .E.......E ....D.V...    50
GII-K74        .....S.... ...N..HQ.. Q....S.L.. .E.......E ....D.I...    50

CI-PFV         RDKFRYLLYT CCATSSRVLA WMFLVCILLI IVLVSCFVTI SRIQWNKDIQ   100
GI-D468        M.RVK.F... .......... ..L.A...F. .II....I.L ..........   100
GII-K74        M.RVK.F... .......... ..L.A...F. .II....I.L ..........   100
                                         |SU
CI-PFV         VLGPVIDWNV TQRAVYQPLQ TRRIARSLRM QHPVPKYVEV NMTSIPQGVY   150
GI-D468        .......... .......... L.....A..A .......... .........F   150
GII-K74        .......... .......... L.....A..A .......... .........F   150

CI-PFV         YEPHPEPIVV KERVLGLSQI LMINSENIAN NANLTQEVKK LLTEMVNEEM   200
GI-D468        .Q......IH T........V .......V.. S...S..T.A .....I....   200
GII-K74        .Q......IH T........V .......V.. S...S..T.A ..........   200
                                        ╟RBD1                |SUvar
CI-PFV         QSLSDVMIDF EIPLGDPRDQ EQYIHRKCYQ EFANCYLVKY KEPKPWPKEG   250
GI-D468        .......... .......... .......... ...H...... .T.Q...S.E   250
GII-K74        .G........ .......... .......... ...H...... .T.Q...N..   250

CI-PFV         LIADQCPLPG YHAGLTYNRQ SIWDYYIKVE SIRPANWTTK SKYGQARLGS   300
GI-D468        .......... ....VE.TT. A......... IT..K...SY AQ..N.....   300
GII-K74        .......... LADVSF.PY. A.....A.I. N.......SS KL..K..M..   300

CI-PFV         FYIPSSLRQI NVSHVLFCSD QLYSKWYNIE NTIEQNERFL LNKLNNLTSG   350
GI-D468        .F..PHV.K- .FT....... ...A...... ..LLK..EL. QK......EL   349
GII-K74        Y...KR..N. .NT.I..... V.......LQ .S.L...NE. TKR.S...-I   349
                                                            ╟RBDj
CI-PFV         TSVLKKRALP KDWSSQGKNA LFREINVLDI CSKPESVILL NTSYYSFSLW   400
GI-D468        ..L....... RT.TT....N ...N.T...V .NR..M.L.. .I..DL....   399
GII-K74        GNK..N.... YE.AKG.L.R ...N.S...V ..R..M.L.. .KT..T....   399

CI-PFV         EGDCNFTKDM ISQLVPECDG FYNNSKWMHM HPYACRFWRS KNEKEETKCR   450
GI-D468        .....Y...K ..EI..Q.K. .......... .........N .........D   449
GII-K74        .....I.RYN VNET....KD .PHRR--FND ...S..L..Y REG...V..L   447
                                                  ╟RBD2    |
CI-PFV         DGETKRCLYY PLWDSPESTY DFGYLAYQKN FPSPICIEQQ KIRDQDYEVY   500
GI-D468        GRDDNK.... .......A.. ...F....N. ..A....SSK Q..Q......   499
GII-K74        TSDHT..... .EYSN..ALF ...F.S.MR. ..G.Q...ST S..Q......   497

CI-PFV         SLYQERKIAS KAYGIDTVLF SLKNFLNYTG TPVNEMPNAR AFVGLIDPKF   550
GI-D468        .I...C.L.. RIH...S... .......... K......... ..........   549
GII-K74        .I...C.L.. .T....S... .......... K......... ..........   547
                   ╢                        |TM
CI-PFV         PPSYPNVTRE HYTSCN--NR KRRSVDNNYA KLRSMGYALT GAVQTLSQIS   598
GI-D468        ..T...I..D Q.QG..INQ. RK.E.N...S .......... ......A...   599
GII-K74        ..T...I..D Q.QG..INQ. RK.E.N...S .......... ......A...   597

CI-PFV         DINDENLQQG IYLLRDHVIT LMEATLHDIS VMEGMFAVQH LHTHLNHLKT   648
GI-D468        ....Q..... .......IV. .......... I......... V.......R.   649
GII-K74        ....Q..... .......IV. .......... I......... V.......R.   647

CI-PFV         MLLERRIDWT YMSSTWLQQ QLQKSDDEMKV IKRIARSLVY YVKQTHSSPT   698
GI-D468        ..M....... ....S...T ........... ...T...... .....YN.L.   699
GII-K74        ..M....... ....S...T ........... ...T...... .....YN.L.   697

CI-PFV         ATAWEIGLYY ELVIPKHIY LNNWNVVNIGH LVKSAGQLTH VTIAHPYEII   748
```

```
GI-D468    .......... ..I..R.... ....Q..... .I........ ..LS......    749
GII-K74    .......... ..I..R.... ....QI.... .I........ ..LS......    747

CI-PFV     NKECVETIYL HLEDCTRQDY VICDVVKIVQ PCGNSSDTSD CPVWAEAVKE    798
GI-D468    .R..SN.L.. ...E.R.L.. .......... .......S.. ......P...    799
GII-K74    .R..SN.L.. ...E.R.L.. .......... .......S.. ......P...    797

CI-PFV     PFVQVNPLKN GSYLVLASST DCQIPPYVPS IVTVNETTSC FGLDFKRPLV    848
GI-D468    .H..IS.... .......... .......... V.......Q. ..VT..K...    849
GII-K74    .H..IS.... .......... .......... V.......Q. ..VT..K...    847

CI-PFV     AEERLSFEPR LPNLQLRLPH LVGIIAKIKG IKIEVTSSGE SIKEQIERAK    898
GI-D468    ...KT.L..Q ..H....... .......... .......... ...D.L....    899
GII-K74    ...KT.L..Q ..H....... .......... .......... ...D.L....    897

CI-PFV     AELLRLDIHE GDTPAWIQQL AAATKDVWPA AASALQGIGN FLSGTAQGIF    948
GI-D468    .......... .......... ....E..... .....K.... ..T.A...L.    949
GII-K74    .......... .......R.. ....E..... .....K.... ..T.A...L.    947

CI-PFV     GTAFSLLGYL KPILIGVGVI LLVILIFKIV SWIPTKKKNQ               988
GI-D468    .....I.... ......I.I. I........L K...I.R.S.               989
GII-K74    .....I.... ......I.I. I........L K...I...S.               987
```

Env sequences from CI-PFV, GI-D468, and GII-K74 strains were aligned using CLC Mainworkbench software. Identical residues are indicated with dots. Boundaries of the leader peptide (LP), surface protein (SU), transmembrane protein (TM), receptor binding domain (RBD)1, RBDj, and RBD2 are indicated over the sequences. The RBDj domain is highlighted by italic characters and the SUvar domain is highlighted by the grey colored background.

#### Supplementary Fig. 2. Recombinant SFV Env oligomerization and mammalian-specific glycolysation do not affect the capacity to inhibit GII-specific nAbs

**A**

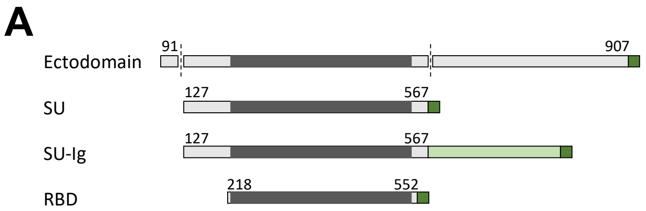

| | GII-K74 | GI-D468 | CI-PFV |
|---|---|---|---|
| Ectodomain | Insect | Not tested | Failed |
| SU | Mammalian<br>Insect | Aggregates | Failed |
| SU-Ig | Mammalian | Aggregates | Mammalian |
| RBD | Mammalian<br>Insect | Not tested | Insect<br>(low level) |

**B**

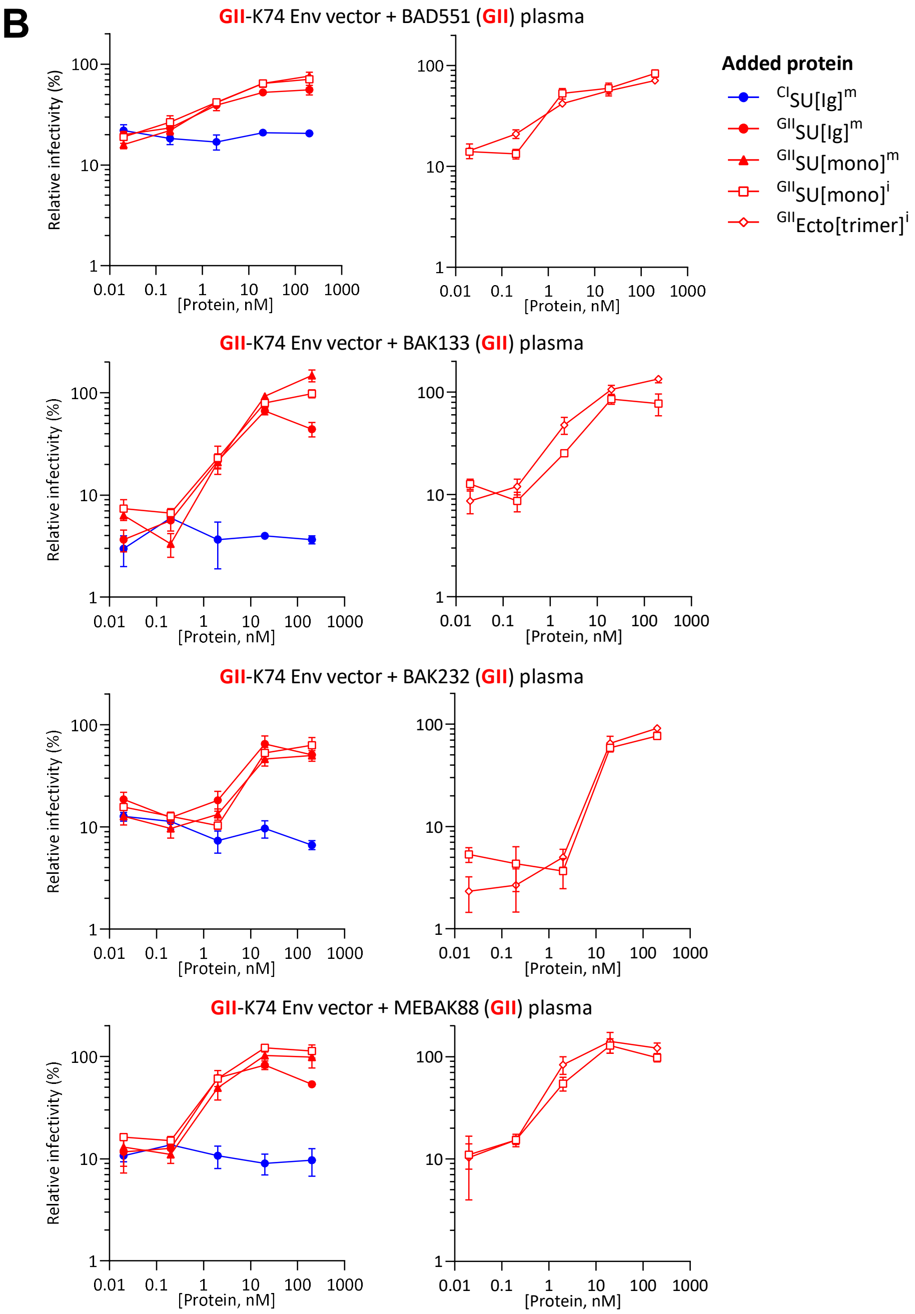

A. Schematic representation of SFV Env constructs tested for expression in mammalian and/or insect cells. SFV Env is shown in grey; the RBD is shwon in dark grey. The dark green segment represents the Twin-Strep-tag and the light green segment the murine Fc domain. The outcomes of expression assays for CI-PFV, GI-D468, and GII-K74 Env-derived proteins are summarized in the table. CI-PFV immunoadhesin ($^{CI}$SU) was the only well-expressed genotype I Env protein and we therefore used immunoadhesins for the project. B. GII-specific plasma samples from four individuals were diluted to their ≈ $IC_{90}$ and incubated with Env-derived proteins at concentrations ranging from 200 to 0.02 nM. The mix was then added to FVVs expressing GII-K74 Env before titration. The relative infectivity is presented as a function of the protein concentration. Production in mammalian or insect cells is indicated in the legend with (m) and (i) suffixes, respectively. The oligomerisation/chimeric state is indicated in parentheses. The left and right panels present independent experiments. The inhibition of anti-GII nAbs by SU was independent of the nature of the producing cell ($^{GII}$SU[mono]$^{m}$ vs. $^{GII}$SU[mono]$^{i}$, left panels), dimerization through fusion with the immunoglobulin constant domain ($^{GII}$SU[mono]$^{m}$ vs. $^{GII}$SU[Ig]$^{m}$, left panels), and trimerization when expressed as an ectodomain in insect cells ($^{GII}$SU[mono]$^{i}$ vs. $^{GII}$Ecto[trimer]$^{i}$, right panels). Genotype-mismatched immunoadhesins ($^{CI}$SU[Ig]$^{m}$) failed to inhibit the nAbs (blue curves, left panels).

#### Supplementary Fig. 3. Western-blot analysis of WT SU proteins used in the study

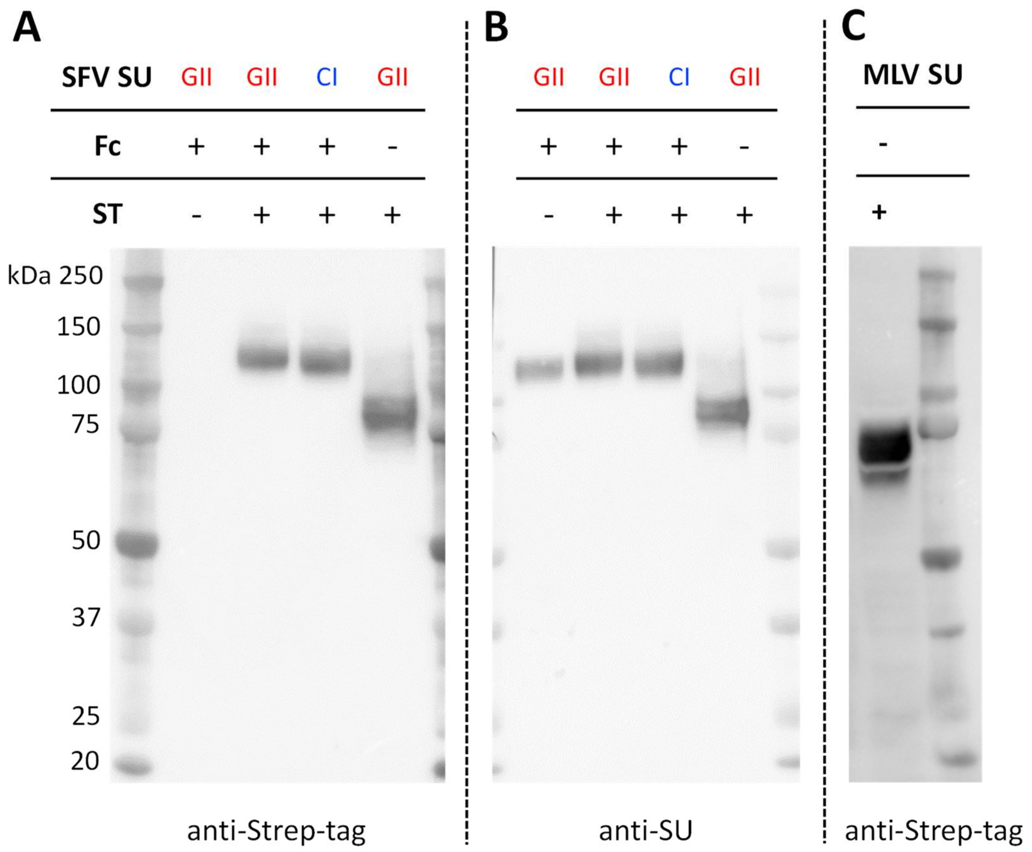

Western-blotting analysis of WT SU protein and immunoadhesins. Mammalian cell supernatants collected 72 h post-transfection were heat-denaturated before immunoblotting with either anti-Strep-tag antibody (A and C) or an anti-SU antibody (B). $^{GII}$SU was expressed as an immunoadhesin without a Strep-tag, an immunoadhesin with a Strep-tag, as monomeric SU with a Strep-tag;s $^{CI}$SU was expressed as an immunoadhesin with a Strep-tag (A and B). $^{MLV}$SU was expressed as SU fused to a Strep-tag (C). For the CI and GII SUs, two bands are visible, in accordance with the results of other reports [27].

#### Supplementary Fig. 4. Purity of proteins used in the study assessed by Coomassie blue gel staining

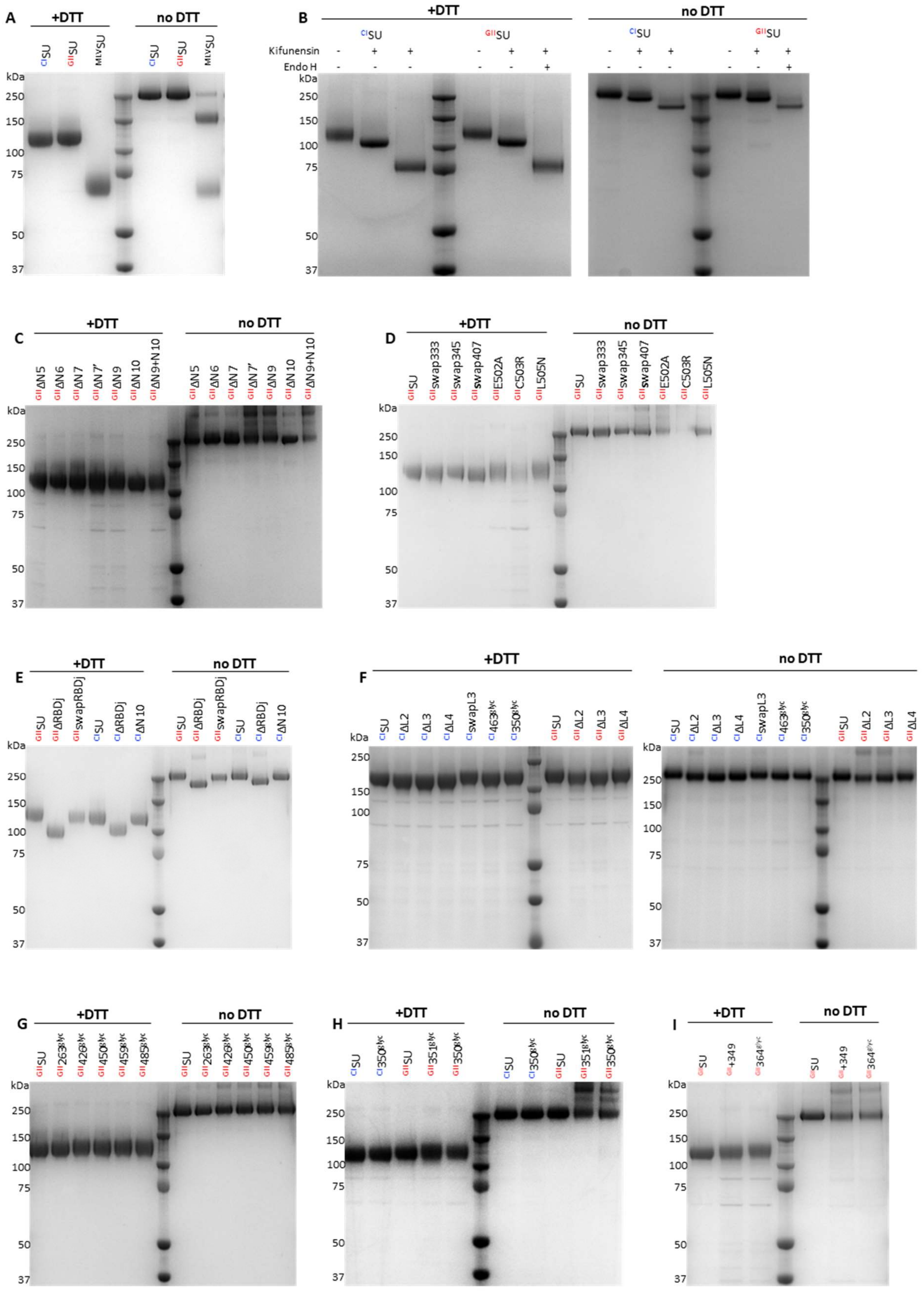

To verify protein purity and aggregate formation, 1.5 µg of purified proteins were heat-denaturated at 70°C for 10 min, with or without DTT. Samples were loaded onto a precast NuPAGE 4-12% Bis-Tris gel and the proteins separated by electrophoresis. Gels were then stained with Coomassie blue and imaged using a G:BOX (Syngene). A western-blot control was performed for all affinity-purified proteins. The constructs are listed in supplementary Table 1 and their names are indicated over the images.

Supplementary Fig. 5. The $^{GII}351_{glyc}$ SU is unable to block nAbs – exclusion of a nonspecific effect of protein aggregation

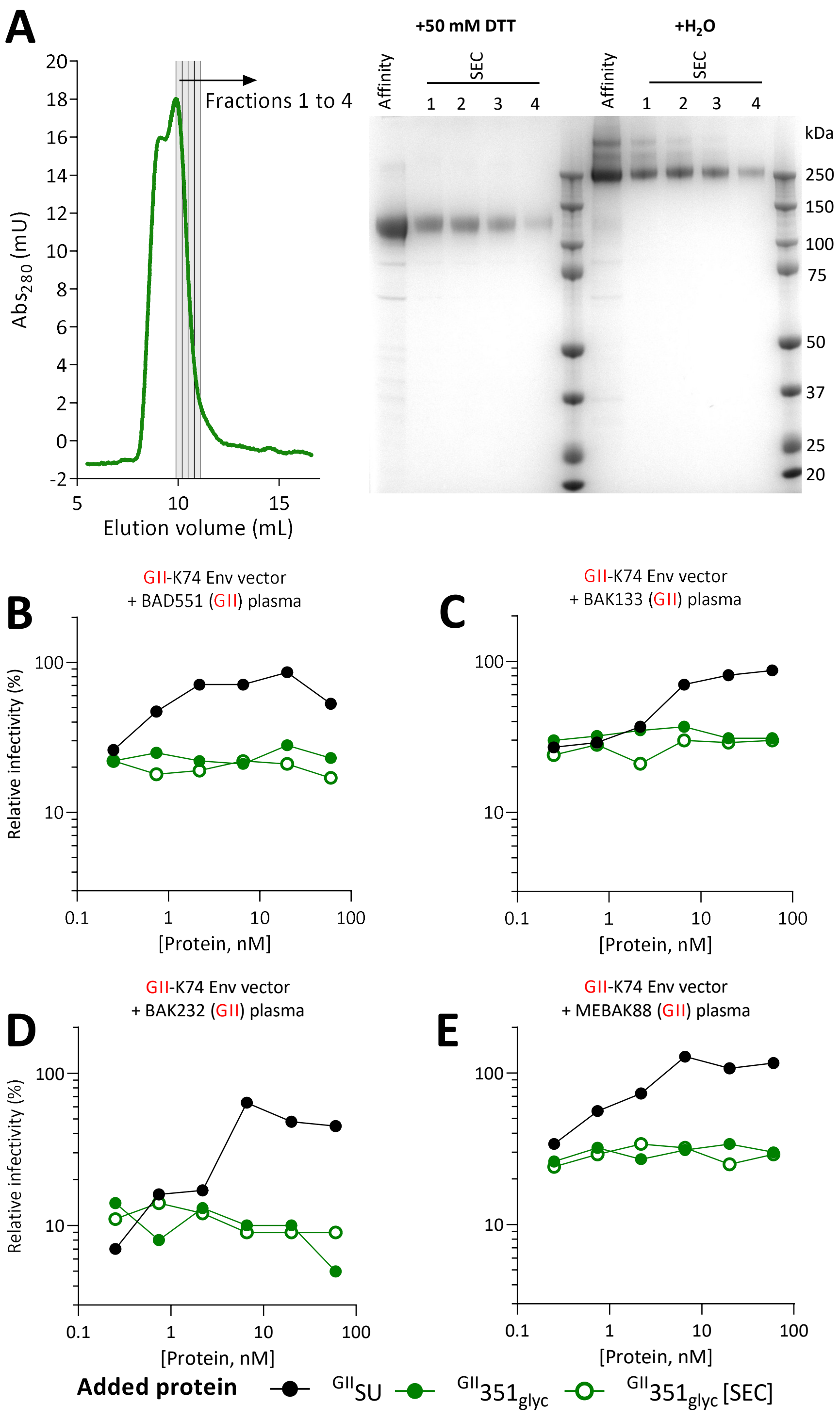

293-F cells were transfected with the plasmid encoding $^{GII}351_{glyc}$. The supernatant was collected after 72 h of culture and the SU was affinity purified using the Strep-Tag fused to the C-terminus. Then, half the volume was purified by size exclusion chromatography. A. The affinity-purified and four fractions of chromatography-purified $^{GII}351_{glyc}$ were analyzed on a Coomassie-stained gel, with or without reducing treatment. High molecular weight proteins were present in the affinity purified sample and the first two SEC fractions. B to E. The plasma samples from four individuals infected with a GII SFV were diluted to their ≈ $IC_{90}$ and incubated with SU at concentrations ranging from 60 to 0.02 nM. The mix was then added to FVVs expressing the GII Env before titrating infectivity. The relative infectivity is presented as a function of SU concentration. The addition of $^{GII}SU$ (black symbols) inhibited the action of the nAbs, whereas affinity-purified $^{GII}351_{glyc}$ did not (green closed symbols). To exclude that $^{GII}351_{glyc}$ aggregation led to epitope masking, the chromatography-purified fractions 3 and 4 ($^{GII}351_{glyc}$ [SEC], green open symbols) were pooled, concentrated, and tested in parallel. These contained no aggregates (panel A) but were unable to block nAbs from the four individuals.

#### Supplementary Fig. 6. The GII chimeric SUs in which aa 333-345 or 345-351 were replaced by GI sequences are unable to block GI-specific nAbs

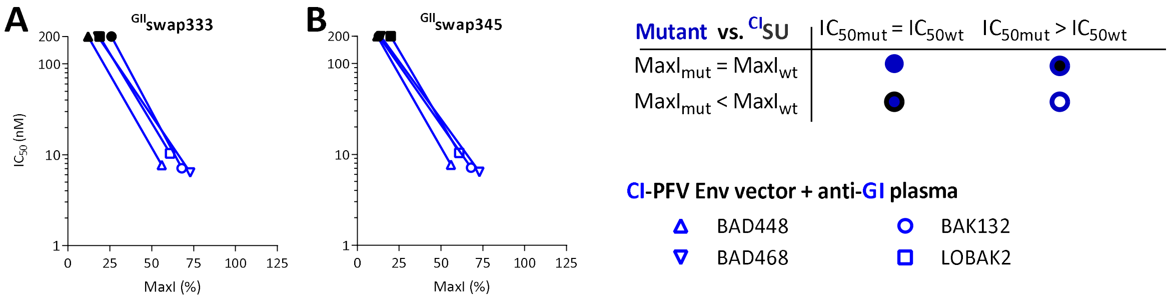

GIISU in which residues 333-345 or 345-351 were replaced by those from the GI-D468 strain were tested for their capacity to block nAbs from GI-specific plasma samples. A. GIIswap333; B. GIIswap345 C. All mutants were tested against four plasma samples. For each plasma sample, the $IC_{50}$ is presented as a function of MaxI for the CISU and mutant SU. The $IC_{50}$ and MaxI values of CISU are presented as open symbols and are those from the same experiment in which the mutant SUs were tested. For mutant SUs, the symbols are colored according to the $IC_{50}$ and MaxI thresholds used to statistically define significant differences from CISU.

#### Supplementary Fig. 7. Only a small proportion of plasma samples bind to peptides covering the SUvar domain

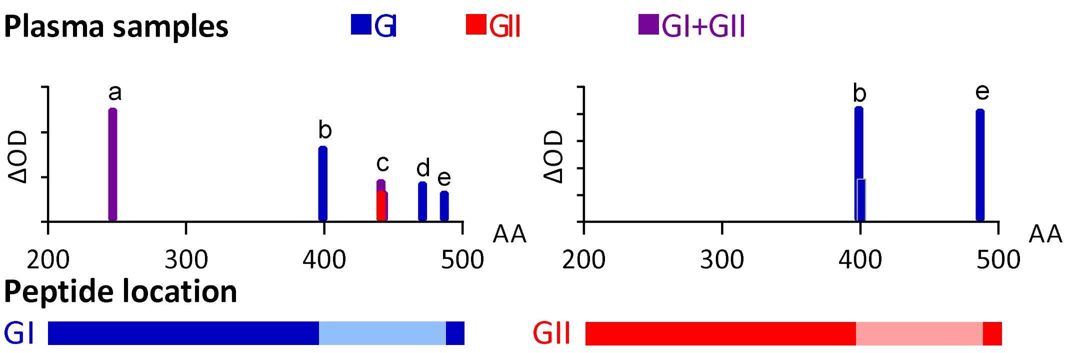

```
CI-PFV      LVKYKEPKPW PKEGLIADQC PLPGYHAGLT YNRQSIWDYY    276
                       BAD468[g1g2]
GI-D468     LVKYKTPQPW PSEELIADQC PLPGYHAGVE YTTQAIWDYY    276

GII-K74     LVKYKTPQPW PNEGLIADQC PLPGLADVSF YPYQAIWDYY    276

IKVESIRPAN WTTKSKYGQA RLGSFYIPSS LRQINVSHVL FCSDQLYSKW    326
IKVEITRPKN WTSYAQYGNA RLGSFFIPPH VRK-NFTHVL FCSDQLYAKW    325
AKIENIRPAN WTSSKLYGKA RMGSYYIPKR LRNINNTHIL FCSDVLYSKW    326

YNIENTIEQN ERFLLNKLNN LTSGTSVLKK RALPKDWSSQ GKNALFREIN    376
YNIENTLLKN EELLQKKLNN LTELTSLLKK RALPRTWTTQ GKNNLFRNIT    375
YNLQNSILQN ENELTKRLSN LT-IGNKLKN RALPYEWAKG GLNRLFRNIS    375

VLDICSKPES VILLNTSYYS FSLWEGDCNF TKDMISQLVP ECDGFYNNSK    426
                         b-BAK56[g1]
VLDVCNRPEM VLLLNISYDL FSLWEGDCNY TKDKISEIVP QCKGFYNNSK    425
                         b-BAK56[g1] BAK82[g1]
VLDVCSRPEM VLLLNKTYYT FSLWEGDCNI TRYNVNETVP ECKDFPHRR-    424

WMHMHPYACR FWRSKNEKEE TKCRDGETKR CLYYPLWDSP ESTYDFGYLA    476
                 c-BAD551[g2] BAK55[g1g2] BAK74[g1g2]
WMHMHPYACR FWRNKNEKEE TKCDGRDDNK CLYYPLWDSP EATYDFGFLA    475
-FNDHPYSCR LWRYREGKEE VKCLTSDHTR CLYYPEYSNP EALFDFGFLS    473

d-BAK132[g1]
YQKNFPSPIC IEQQKIRDQD YEVYSLYQER KIASKAYGID TVLFSLKNFL    526
          e-BAK56[g1]
YQNNFPAPIC ISSKQIRQQD YEVYSIYQEC KLASRIHGID SVLFSLKNFL    525
          e-BAK56[g1]
YMRNFPGPQC IESTSIRQQD YEVYSIYQEC KLASKTYGID SVLFSLKNFL    523
```

Seventeen plasma samples from African hunters were tested for binding to 37 peptides located in the SUvar domain (Supplementary Table 5). The summary graph shows positive responses ($\Delta_{OD}$, y-axis) plotted against peptides identified by the position of their first aa (x-axis). Plasma samples are identified by a color code corresponding to the genotype(s) of the infecting strains (blue: infected with a GI strain, red: infected with a GII strain, purple: infected by strains of both genotypes [7]). Left, peptides spanning GI SUvar; right, peptides spanning GII SUvar; the RBDj region is indicated by the lighter color. Detailed binding activity is shown on the aligned SU sequences; the RBDj sequence is highlighted in italic characters; the sequence covered by peptides is highlighted by grey background. Recognized peptides are underscored and designated by the same letters as those used in panel A. Reactive plasma samples are indicated above the sequence and colored according to the genotype(s) of the infecting strains. Six plasma samples from SFV-infected individuals reacted against seven peptides located in the RBDj region (BAD551, BAK55, BAK56, BAK74, BAK82, and BAK132) and two samples reacted against a peptide located in the RBD1 or RBD2 subdomains (BAD468 and BAK56, respectively). Plasma antibody binding to the peptides was not genotype-specific. For example, sample BAK56 reacted against peptide b (399W-K418) from the GI-D468 and GII-K74 strains.

#### Supplementary Fig. 8. Recombinant SU bind to susceptible cells

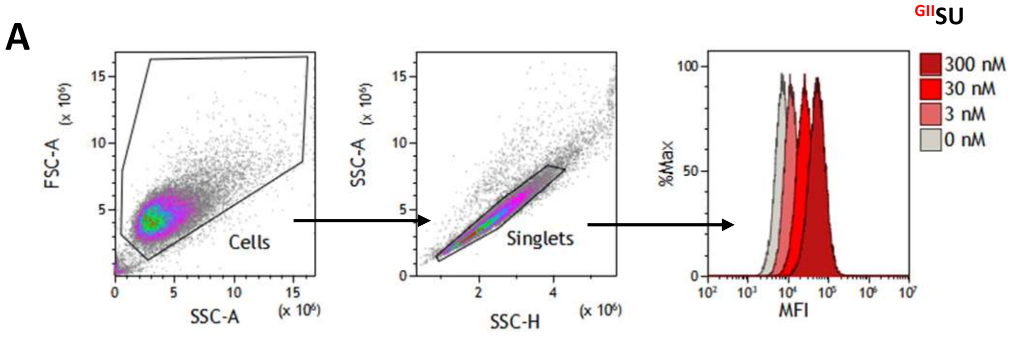

| Concentration | MFI | MFIprotein/untreated |
|---|---|---|
| 300 | 38113 | 5.11 |
| 30 | 18977 | 2.54 |
| 3 | 12699 | 1.70 |
| 0 | 7462 | - |

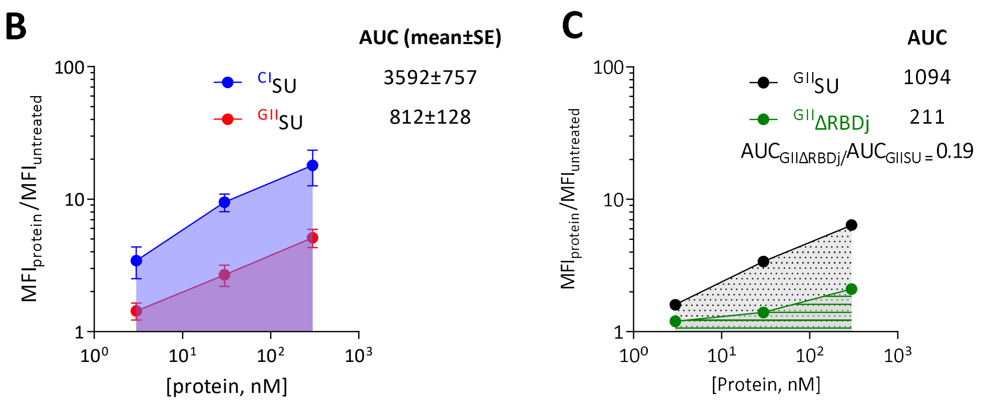

HT1080 cells were incubated with SU and the bound protein detected by staining with an anti-mouse Fc antibody and flow cytometry analysis. A. The gating strategy of viable single cells and staining intensity are shown for $^{GII}$SU added at three concentrations. Levels of bound $^{GII}$SU are expressed as the ratio of the MFI from SU-treated cells to the MFI of untreated cells. B. To compare the binding capacity of the SU, staining was performed at three doses, the MFI ratios plotted as a function of SU concentration, and the area under the curve (AUC) calculated. Shaded regions represent the AUC. Data from five independent experiments performed with WT SU are presented as the mean and standard error. $^{CI}$SU bound at higher levels than $^{GII}$SU. C. The treated and mutated SU were tested for binding to susceptible cells and staining levels were normalized to that of the WT SU included in every experiment. The graph shows lower staining by $^{GII}\Delta$RBDj than $^{GII}$SU.

Supplementary Fig. 9. Sequences from gorilla SFV strains circulating in Central Africa are conserved in the epitopic regions targeted by nAbs

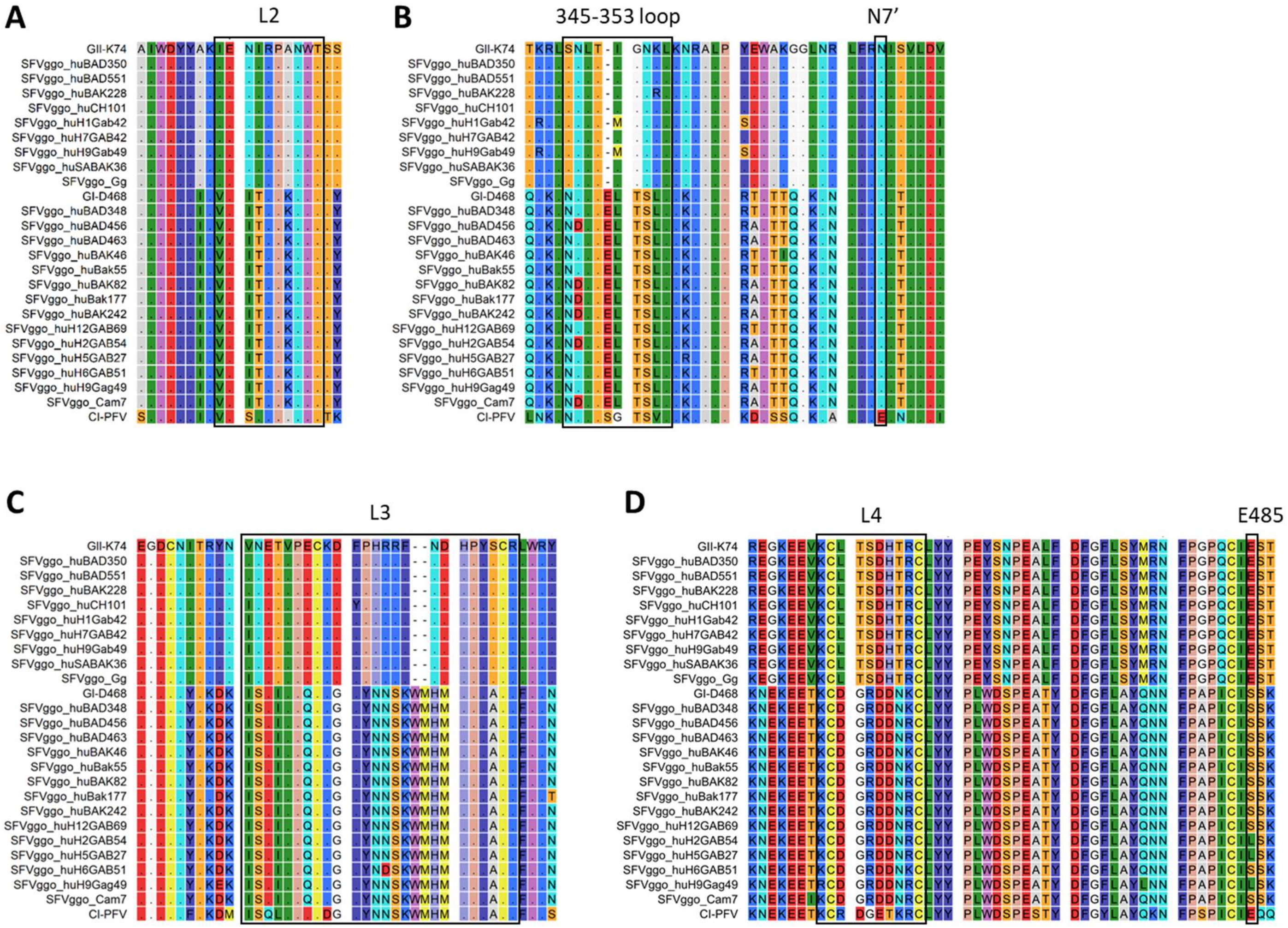

The SUvar protein sequences from the SFVggo strains circulating in Central Africa and CI-PFV were aligned. We included all available sequences: 10 genotype II sequences (nine zoonotic strains and one animal strain [11]) and 15 genotype I sequences (14 zoonotic strains and one animal strain [11]). Identical residues are indicated by dots, the background colors correspond to the physical properties (rasmol color code). Black squares indicate the identified epitopic regions: L2 (A), 345-353 loop and N7' (B), L3 (C), L4 and E485 (D). Within each genotype, we observed identical sequences or conservative aa changes. The only exception was an N351/D polymorphism in the 345-353 loop from GI that may alter the expression of N7 strain.
